# Spatial transcriptomics of *C. elegans* males and hermaphrodites identifies novel fertility genes

**DOI:** 10.1101/348201

**Authors:** Annabel Ebbing, Abel Vertesy, Marco Betist, Bastiaan Spanjaard, Jan Philipp Junker, Eugene Berezikov, Alexander van Oudenaarden, Hendrik C. Korswagen

## Abstract

To advance our understanding of the genetic programs that drive cell and tissue specialization, it is necessary to obtain a comprehensive overview of gene expression patterns. Here, we have used RNA tomography to generate the first high-resolution, anteroposterior gene expression maps of *C. elegans* males and hermaphrodites. To explore these maps, we have developed computational methods for discovering region and tissue-specific genes. Moreover, by combining pattern-based analysis with differential gene expression analysis, we have found extensive sex-specific gene expression differences in the germline and sperm. We have also identified genes that are specifically expressed in the male reproductive tract, including a group of uncharacterized genes that encode small secreted proteins that are required for male fertility. We conclude that spatial gene expression maps provide a powerful resource for identifying novel tissue-specific gene functions in *C. elegans*. Importantly, we found that expression maps from different animals can be precisely aligned, which opens up new possibilities for transcriptome-wide comparisons of gene expression patterns.

## Introduction

Examination of spatial gene expression patterns provides important insight into the gene expression programs that drive the functional specialization of cells and tissues in multicellular organisms. A model organism that is ideally suited to connect such spatial expression information to cellular functions is the nematode *C. elegans*. It has a relatively simple body plan of 959 somatic cells in adult hermaphrodites and 1031 somatic cells in adult males (Sulston et al., 1980; Sulston and Horvitz, 1977), which enables gene expression pattern analysis at cellular resolution in a complete organism. Moreover, there is considerable complexity in specialized cell and tissue types (Sulston and Horvitz, 1977) to link expression patterns to specific evolutionarily conserved functions.

Gene expression analysis in *C. elegans* has mostly relied on microscopy-based techniques such as mRNA *in situ* hybridization, immunohistochemistry and promoter analysis with reporter transgenes. Even though large-scale *in situ* hybridization and promoter studies have been performed (Dupuy et al., 2007; Tabara et al., 1996), a transcriptome-wide overview of gene expression patterns has remained out of reach. Moreover, since most gene expression studies have focused on the hermaphrodite, expression patterns in the male are still largely uncharacterized (Kim et al., 2016).

RNA sequencing is a powerful approach to study gene expression at the whole-genome level, but it does not provide spatial information. Gene expression analysis on isolated cell populations partially solves this problem, but isolation of certain cell types can be problematic and spatial information is limited (Cao et al., 2017). An approach to overcome these limitations is to combine RNA sequencing with serial tissue-sectioning (Junker et al., 2014). Here, spatial information is retained by cryo-sectioning the sample along a specific axis, and then performing RNA sequencing on the individual sections. This method, which is referred to as RNA tomography or tomo-sequencing (tomo-seq), has been used to create a three-dimensional gene expression atlas of the zebrafish embryo (Junker et al., 2014) and has also been applied to study expression patterns in organs such as the heart (Wu et al., 2016).

A major challenge in applying RNA tomography to small organisms such as *C. elegans* is the limited amount of mRNA that can be extracted from individual sections, especially when thin sections are used to create high-resolution gene expression maps. Here, we have used the sensitive CEL-seq method (Hashimshony et al., 2012) to create the first high-resolution, anteroposterior (AP) gene expression maps of *C. elegans* males and hermaphrodites. We have developed computational methods to align and cluster expression maps into distinct anatomical regions and to identify spatially co-expressed genes based on the similarity of expression patterns. Using this approach, we have identified genes specific to the germline, sperm, and different somatic cells and tissues. Furthermore, by combining these results with sex-specific differential gene expression analysis, we have identified male and hermaphrodite specific gene expression differences in the germline and in sperm. Using a similar strategy, we discovered somatic genes specific to the male reproductive system, including a novel set of reproductive tract genes encoding secreted proteins that are required for male fertility. Our results demonstrate that RNA tomography maps provide a powerful resource to identify novel sex- and tissue-specific gene functions in *C. elegans*.

## Results

### Generation of high-resolution, transcriptome-wide anteroposterior gene expression maps of *C. elegans*

To generate high-resolution gene expression maps of *C. elegans*, we adapted the RNA tomography approach developed for zebrafish (Junker et al., 2014). In brief, we individually froze and cryo-sectioned young adult male and hermaphrodite animals into 20 μm thick sections along the anteroposterior (AP) body axis (Fig. 1A). RNA was extracted from each section using Trizol and processed using a unique molecular identifier (UMI) extended CEL-seq protocol (Grun et al., 2014). In this approach, each detected molecule is labeled by a UMI (Kivioja et al., 2011) and a section specific barcode before the samples are pooled to prepare a single Illumina sequencing library. The resulting sequencing reads were trimmed for low quality bases and aligned to the *C. elegans* reference transcriptome (WS249) by BWA-MEM. Finally, the spatial distribution of the transcripts was recovered using the section specific barcodes.

We sectioned and analyzed 4 young adult hermaphrodites and 4 young adult males. In addition, we included 2 germline deficient *glp-1(q231)* hermaphrodites (Austin and Kimble, 1987) to examine somatic gene expression in the mid-body without interference from the mRNA-rich germline. Despite the low amount of input material (sections that do not cover the germline contain on average only about 20-30 somatic cells), our approach yielded high-complexity data. After optimizing sequencing depth, we obtained 13-22 million uniquely mapped, unduplicated reads, corresponding to 4-11 million unique transcripts per sectioned wild type male or hermaphrodite animal (Fig. 1B, Table S1). We could detect an average of 16394 genes in hermaphrodites and 16994 in males, with a total 18778 and 19241 genes in the combined datasets, respectively. This represents 93% and 95.3% of genes that were detected in a recent study using bulk RNA sequencing of young adult hermaphrodites and males (Kim et al., 2016).

**Figure 1.**
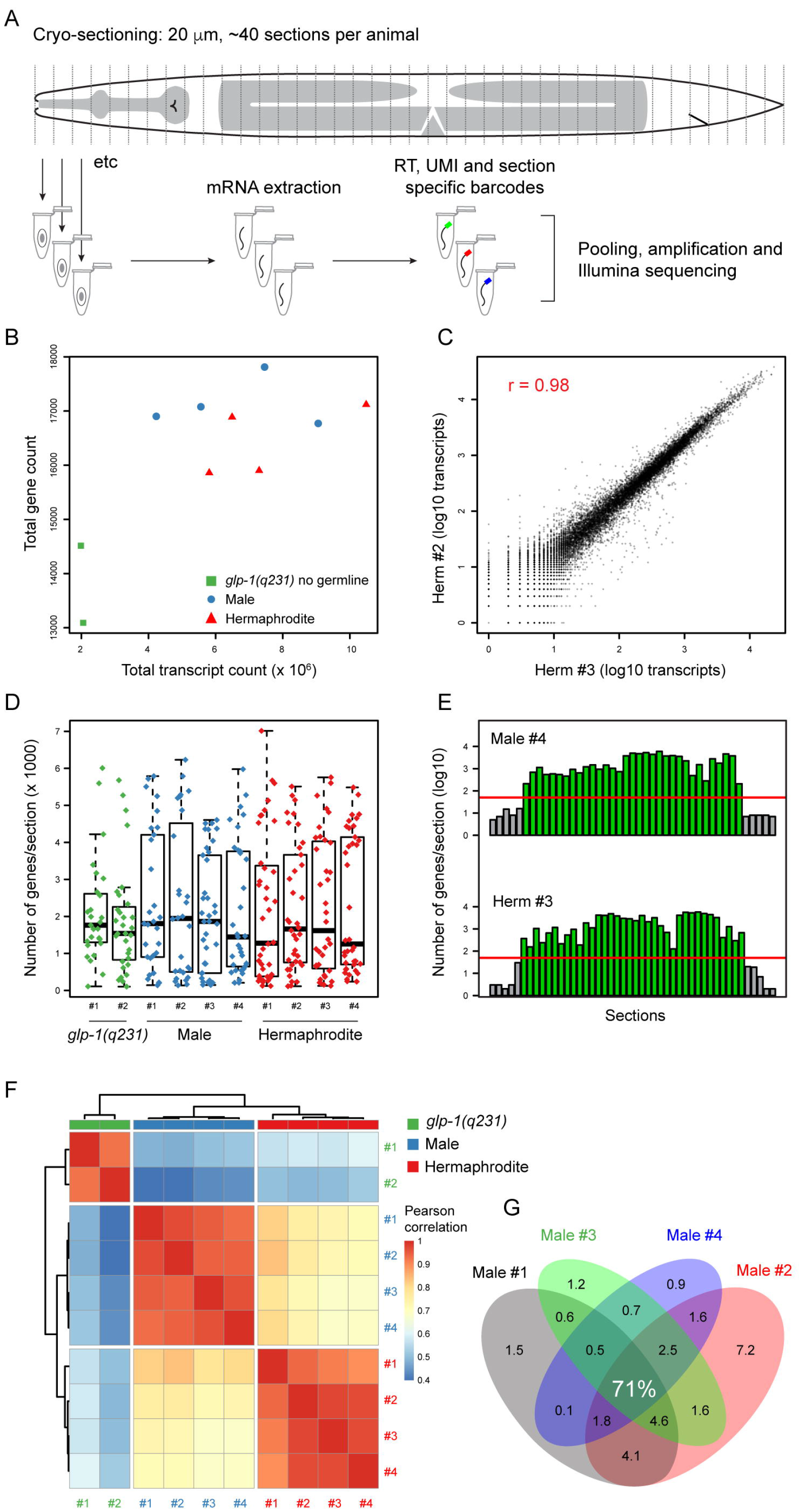
RNA tomography in *C. elegans*. (**A**) Schematic overview of the tomo-sequencing method. Individual young adult animals were sectioned along the AP axis into 30-45, 20 μm thick sections. Each section underwent mRNA extraction, followed by reverse transcription and addition of section specific barcodes and unique molecular identifiers (UMIs). Next, the samples were pooled and subjected to linear amplification by *in vitro* transcription, followed by reverse transcription, library preparation and Illumina sequencing. (**B**) Total number of genes and transcripts per young adult hermaphrodite and male dataset. The 2 young adult *glp-1(q231)* germline deficient mutants were sequenced at lower depth. (**C**) Pearson correlation (r) of total transcript data in Hermaphrodite datasets #2 and #3. (**D**) Total number of genes detected. Mean and standard deviation are indicated. (**E**) Distribution of the number of genes detected per section along the AP axis. Red line indicates threshold of 50 genes per section. (**F**) Pearson correlation clustering segregates the datasets by sex and shows high reproducibility across datasets of the same sex. (**G**) Venn diagram of genes with >10 transcripts in at least 1 section in the 4 male datasets. >70% of genes are detected in each of the 4 datasets.

For further analysis, we kept all genes expressed above 10 unique transcripts in at least one section. The majority of these genes were detected in all datasets of the same sex (71% in males and 69% in hermaphrodites, Fig. 1G, S1A). Sequencing libraries were normalized to 10 million total transcripts per animal. Next, the anterior and posterior ends of the expression maps were defined by the first or last two consecutive sections with ≥50 genes (with ≥20 transcripts each). Internal sections with <50 genes were removed (an average of 1.1 sections per animal). The total number of high-quality sections per animal was, on average, 41 sections for the hermaphrodite and 34 for the male datasets, which is consistent with the difference in length between the two sexes. Along the sectioning axis, there was an average of 2170 genes per section in hermaphrodites and 2210 in males (Fig. 1D, E). Interestingly, we found that 39 genes show clear spatial differences in 3’ isoform usage, most notably between the soma and germline, or between different regions of the germline (Fig. S1B, Table S2, Supplementary dataset 1). Sections within the boundaries of the animal were z-score transformed, or normalized to 10 million reads per animal, or converted to transcript per million (TPM) values per section for specific analyses. When comparing the different hermaphrodite and male datasets, Pearson correlation coefficients of the pooled transcriptome data ranged from 0.90-0.98 for hermaphrodites and 0.95-0.97 for males, demonstrating the high reproducibility of the method (Fig. 1C). As expected, transcriptome-wide Pearson correlation distinguished the male and hermaphrodite datasets by sexual identity (Fig. 1F). Taken together, these results show that RNA tomography is a sensitive and reproducible method to examine gene expression in *C. elegans*.

### Expression patterns of marker genes validate the high spatial resolution and sensitivity of *C. elegans* RNA tomography maps

To get a global view of gene expression patterns along the AP axis, we performed hierarchal clustering on the z-score normalized gene expression data. As shown in Fig. 2A and B, several large clusters of co-expressed genes could be identified in males and hermaphrodites: an anterior cluster including genes known to be specifically expressed in the head region, several clusters in the mid-body region that represent the germline and sperm and a group of genes in the posterior that includes tail specific genes. To examine these expression patterns in more detail, we plotted the expression of marker genes that are specific to major anatomical structures such as the pharynx, the nerve ring, the germline, the spermatheca and vulva, as well as the male reproductive tract and tail (Fig. 2C, E). As expected, the expression patterns of these marker genes were consistent with the anatomical localization and spacing of the corresponding structures along the anteroposterior body axis in the intact animal. Importantly, also genes expressed in specific pairs of neurons could be readily detected, demonstrating the high sensitivity of our approach. An example is *cwp-1*, which is expressed in only two adjacent neuron pairs in the head (the male specific ventral CEMV and dorsal CEMD neurons) (Portman and Emmons, 2004), and shows the expected single peak of expression in the anterior region of the male expression maps (Fig. 2F). Finally, we found that the spatial resolution of the expression patterns corresponded well with the 20 μm sectioning width. For example, the IL1, OLL and URB neurons that express the neuropeptide gene *flp-3* (Li et al., 1999) and neurons of the nerve ring that express *flp-1* (Nelson et al., 1998) are positioned approximately 60 μm apart in the head region. As expected, we found that the peaks of *flp-3* and *flp-1* expression were 2-3 sections apart in the hermaphrodite expression maps (Fig. 2D).

**Figure 2.**
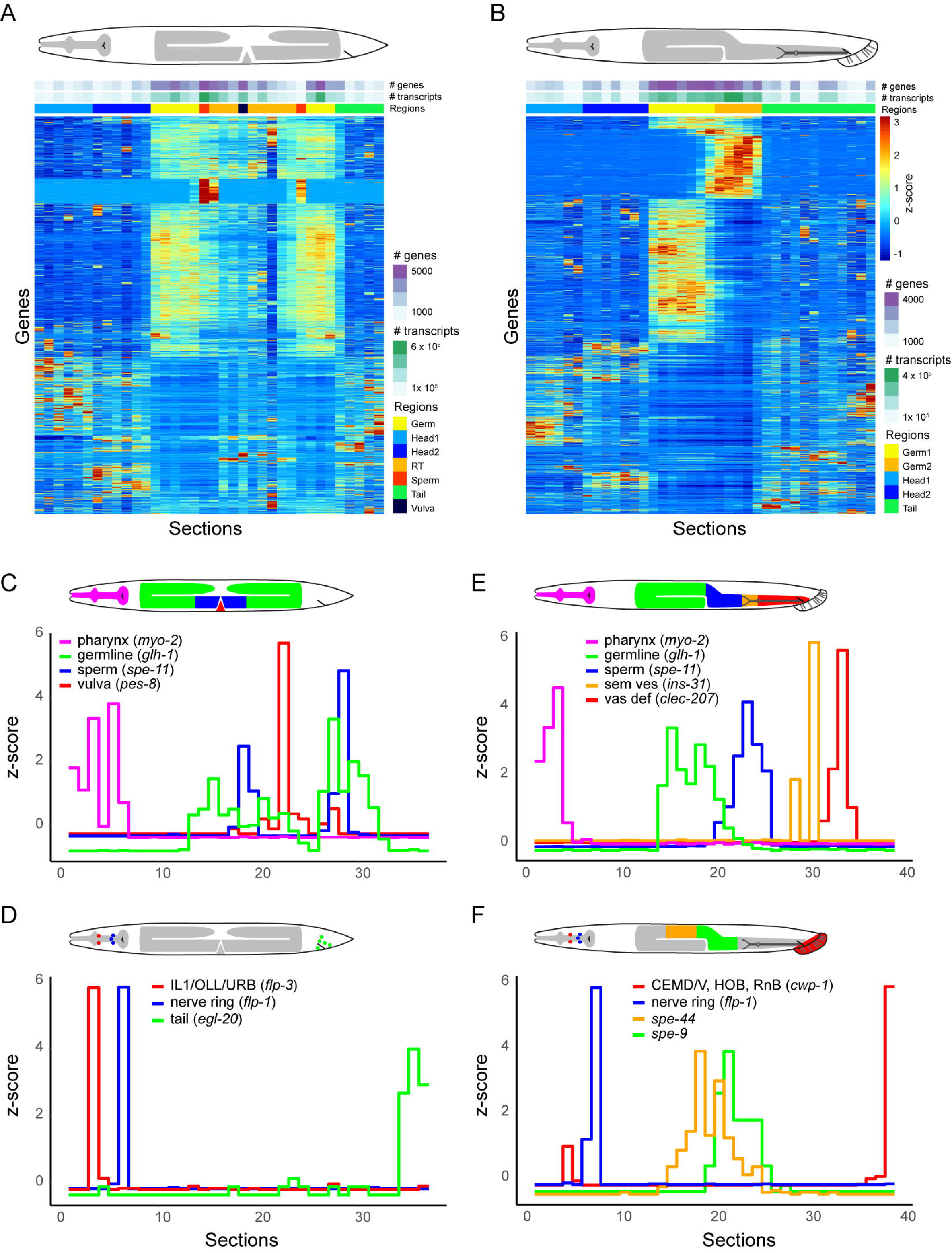
Validation of RNA tomography expression maps. (**A**, **B**) Transcript distribution correlates with anatomical structures. Heatmaps of z-score normalized gene expression in Hermaphrodite #3 and Male #3 (genes with ≥ 15 normalized reads in ≥2 consecutive sections are shown). Pearson correlation based clustering shows groups of genes specific to the head, germline and tail. The largest set of genes in hermaphrodites is germline related and is expressed in a symmetric way around the vulva. Anatomical features such as the head, germline (Germ), hermaphrodite reproductive tract (RT), vulva, sperm and tail are indicated. (**C**, **D**) Expression of representative cell and tissue-specific marker genes in hermaphrodites and (**E**, **F**) males. The symmetric hermaphrodite germline anatomy (green) and the linear spermatogenic and reproductive region (blue, orange, red) in males are clearly delineated. The AP expression of *spe-44* (initiation of spermatogenesis) and *spe-9* (mature spermatids) correlates with the different stages of male spermatogenesis. Note that the colour coded drawings of the expression patterns are not at the same scale as the graphs.

### Unbiased computational and manually curated alignment of male and hermaphrodite expression maps identifies region-specific genes

Although the order of gene expression patterns along the anteroposterior axis was highly consistent between expression maps of different animals, we found some variability in the spacing of expression peaks (Fig. S2). This is most likely caused by subtle differences in the length of the animals and technical variability of the cryo-sectioning procedure. However, since the anatomical positions of cells in *C. elegans* are largely invariable (Sulston et al., 1980; Sulston and Horvitz, 1977), we could align and merge expression maps using (1) unbiased computational clustering and (2) manually-curated alignment.

In the first approach, we assigned each section to a distinct region by sequentially breaking down the expression map into smaller regions using hierarchical clustering. In brief, we first calculated pairwise Pearson correlations between all sections using the TPM-normalized data. Male sections were first separated into the 3 most distinct clusters, resulting in an anterior germline region (Germ1), a posterior germline region (Germ2) and a somatic cluster (see supplemental experimental procedures for detailed description of the clustering approach). The Germ1 cluster corresponds to the distal part of the germline, while Germ2 corresponds to the spermatogenic region of the germline. The somatic cluster contained the head and tail regions, which could be separated by their AP position. Moreover, we found that the head region could be further divided into two sub-clusters (Head1 and Head2), which correspond to the anterior part of the head, and the nerve ring plus anterior intestine, respectively (Fig. 3A). As hermaphrodites have a more complex gonadal structure (two symmetric gonad arms centered around the uterus and vulva), we applied a hybrid approach combining clustering with the use of marker genes. First, we defined the vulva as the section with the highest expression of the marker gene *pes-8* (Hope et al., 1998) (Fig S3A). Next, we defined the two regions where sperm is stored as anterior and posterior expression maxima of the marker gene *msp-3* (Kosinski et al., 2005). These 3 sections separate the animal into 4 regions: an anterior region, two reproductive tract regions RT.a and RT.p flanking the vulva, and a posterior region. The anterior region was then separated into the three most distinct sub-clusters: Head1 (pharynx and nerve ring), Head2 (anterior intestine) and Germ.a (anterior arm of the gonad); and the posterior region was divided into two sub-clusters: Germ.p and Tail (Fig. 3B). In agreement with the symmetric structure of the hermaphrodite germline, the Germ.a and Germ.p regions showed high pairwise correlation with each other. Importantly, we found that a similar hierarchical clustering strategy could be used to align and merge sections from all sequenced animals of the same sex. As shown in Fig. 3C for the 4 male datasets (and Fig. S3B for the 4 hermaphrodite datasets), we found that sections separated by the same AP regions as in the individual animals, but not by sample of origin.

**Figure 3.**
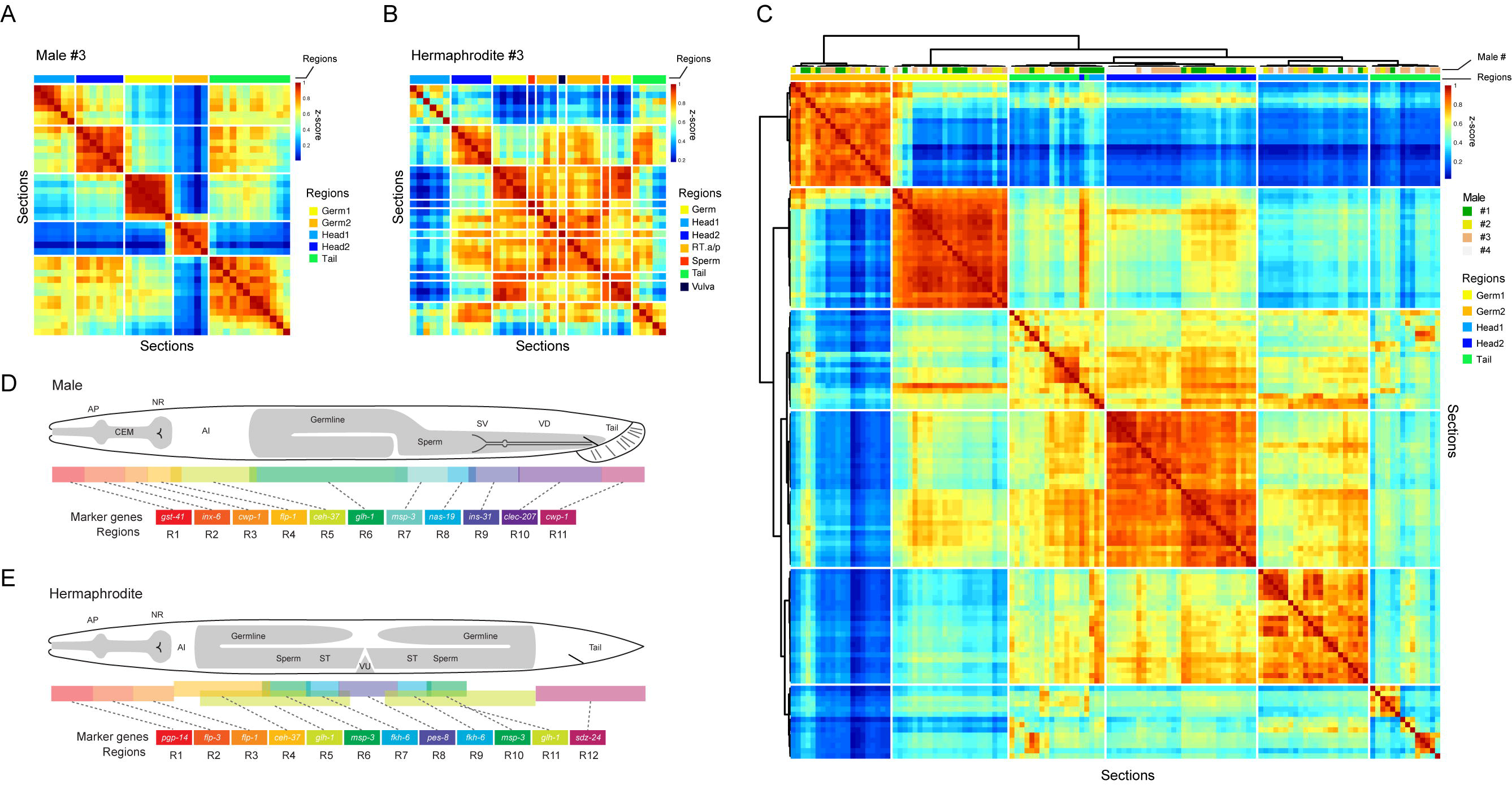
Transcriptome-wide clustering separates major anatomical regions along the AP axis. (**A**) Pairwise Pearson correlation heatmap of Male #3 shows distinct gene expression in 5 major anatomical regions. Column annotation shows the 5 regions identified by sequential hierarchical clustering. (**B**) Pairwise Pearson correlation heatmap of Hermaphrodite #3 shows the symmetric germline architecture. (**C**) Major regions cluster by anatomical identity rather than sample of origin when all sections from all four male datasets are clustered together. (**D**, **E**) Complementary manually curated annotation based on marker gene expression identifies specific regions at a higher resolution. The datasets were aligned and merged based on these markers. AP anterior pharynx region, NR nerve ring, AI anterior intestine region, ST spermatheca, VU vulva and uterus region, SV seminal vesicle, VD vas deferens.

In the second approach, we manually aligned the different expression maps using marker genes showing a unique expression peak at key locations along the AP body axis and merged the remaining sections with their most similar neighbors (see supplemental experimental procedures for details). Next, we combined the corresponding regions for each sex, and calculated the average normalized expression. This approach yielded merged expression maps that are of higher resolution than could be obtained with hierarchical clustering: the hermaphrodite expression maps could be divided into 12 regions that include different sections of the head, the anterior intestine, vulva and tail, and the symmetrical structure of the germline and reproductive system (Fig. 3E, S3C). In males, 11 regions were defined that correspond to different parts of the head, the anterior intestine, the germline, parts of the reproductive tract and the tail (Fig. 3D, S3D).

An important advantage of the merged expression maps is an increase in sensitivity and a corresponding decrease in noise. The merged expression maps are therefore especially useful for examining genes that are expressed at a low level, which can subsequently be further investigated in the individual high-resolution datasets. Cluster heatmaps of all male and hermaphrodite datasets, and comparisons of clustering to the marker genes used in the manual alignment are provided (Supplementary dataset 2). To facilitate the discovery of tissue-specific genes, the expression pattern of each gene (per section and per manually assigned region) can be viewed interactively on the project website (http://celegans.tomoseq.genomes.nl).

### Identification of tissue-specific genes based on expression pattern similarity

Each gene in the tomo-seq expression maps displays a specific pattern along the AP axis. A germline gene, for example, shows a complex series of peaks that correspond to the anterior and posterior arms of the hermaphrodite gonad, while a neuron specific gene such as *flp-3*, shows only a single peak in the head region (Fig. 2C, D). The peak pattern of a gene, therefore, provides a unique fingerprint that can be used to identify other genes with a similar expression pattern. We developed a computational approach to identify co-expressed genes using Pearson correlation of z-score normalized gene expression across sections (see supplemental experimental procedures for details). Using this algorithm, the germline genes *glh-1* and *pgl-1*, for example, show a Pearson correlation coefficient (r) of 0.98, while the correlation coefficient of *glh-1* with the ubiquitously expressed household gene *act-*1/actin is only 0.68 (Fig. 4A, B). Thus, co-expression of genes can be reliably detected, even when their expression patterns are complex.

**Figure 4.**
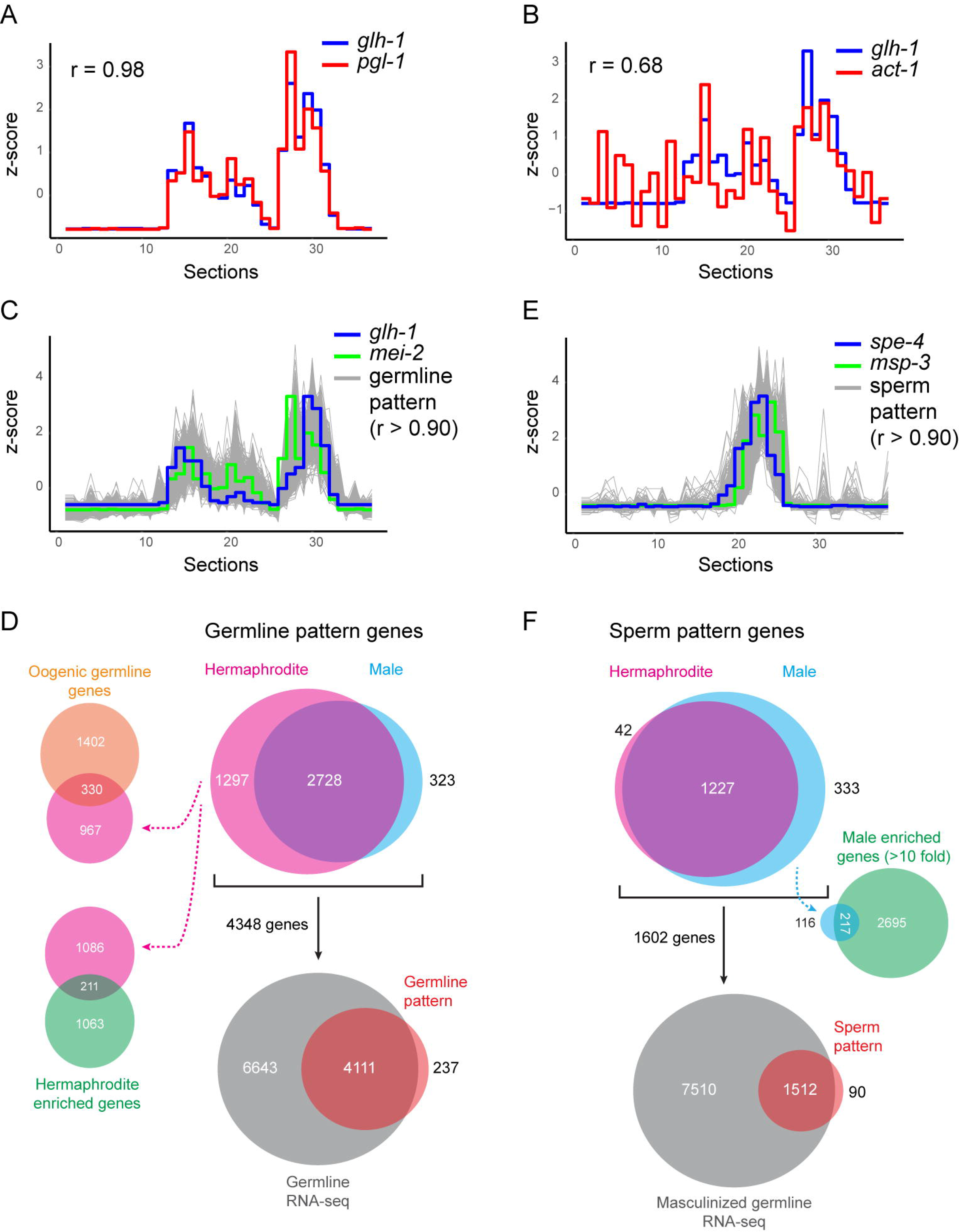
Identification of germline and sperm specific genes using spatial co-expression. (**A**, **B**) Quantification of spatial co-expression using Pearson correlation. Examples of coexpression between the germline genes *glh-1* and *pgl-1* and between *glh-1* and the ubiquitously expressed gene *act-1* in Hermaphrodite #3. (**C**) 4025 germline specific genes were identified in hermaphrodites using co-expression similarity to *glh-1* or *mei-2* at a Pearson correlation value > 0.90. (**D**) 94.5% of the 4348 combined hermaphrodite and male germline pattern genes overlap with genes identified by bulk RNA sequencing of dissected gonads (Ortiz et al., 2014). 2728 germline pattern genes are shared between hermaphrodites and males, while 1297 were only identified in hermaphrodites. 211 of these genes are 2-fold enriched in hermaphrodites and 330 overlap with bulk RNA sequencing data of oogenic germlines (Ortiz et al., 2014). (**E**) 1560 sperm specific genes were identified in males using co-expression similarity to *spe-4* or *msp-3* at a Pearson correlation value > 0.90. Expression in Male #3 is shown. (**F**) 94.4% of the 1602 combined hermaphrodite and male sperm pattern genes overlap with genes identified by bulk RNA sequencing of dissected masculinized gonads (Ortiz et al., 2014). 1227 sperm pattern genes are shared between males and hermaphrodites. 217 of the 333 sperm pattern genes only detected in males are >10-fold enriched in the male transcriptome.

We found that peak pattern similarity provides a powerful approach to identify tissue-specific genes. By taking a gene with a known expression pattern as an anchor point and setting a stringent threshold, this approach identifies other genes in the dataset that show a highly similar expression pattern. These we call *“pattern specific genes”*. An example is the identification of sperm pattern genes using the sperm genes *spe-4* and *msp-3* as anchor genes (Fig. 4E). Using a threshold of r > 0.90, we found a total of 1602 genes in males and hermaphrodites. These include 30 out of 47 sperm specific *msp* genes and 19 out of 26 *spe* genes annotated in Wormbase (version WS260) (Table S3). Furthermore, there is 94% overlap with genes expressed in dissected masculinized germlines (Fig. 4F) (Ortiz et al., 2014). These results indicate that we have identified the majority of sperm specific genes and validate our approach to discover tissue-specific genes. Importantly, because of the stringent criteria for similarity, our approach enriches for genes that are specific to the tissue. This is an advantage over RNA sequencing of isolated cells or tissues, which detects all genes, including housekeeping genes that are expressed throughout the organism.

Because our approach is based on similarity in peak patterns, there is little overlap between genes identified with different anchor genes, even when these are expressed in close proximity. The seminal vesicle and vas deferens, for example, are located next to each other in the male reproductive tract (Fig. 2E, 5D, 5G), but only show 7 overlapping genes out of 125 seminal vesicle and 323 vas deferens specific pattern genes, respectively (Table S3). Encouraged by these results, we identified genes with specific expression patterns for a range of tissues and organs, including the anterior head region, the nerve ring, the pharynx, the germline and sperm, the spermatheca, the vulva and uterus, the male CEM neurons and the male reproductive tract and tail (Table S3). Furthermore, an interactive search tool is available on the project website.

### Spatial gene expression analysis identifies sexually dimorphic germline and sperm-specific genes

The generation of both hermaphrodite and male expression maps allowed us to compare sex-specific gene expression in different tissues and organs. A tissue of particular interest is the germline. In adult hermaphrodites, the germline produces oocytes that are loaded with the maternal products that are required for the early stages of embryogenesis, while in males the germline produces sperm (Bowerman, 1998; Ellis and Stanfield, 2014; Kimble and Crittenden, 2007; Sulston et al., 1980). To identify germline specific genes in hermaphrodites and males, we used our pattern similarity algorithm to search for genes co-expressed (at r > 0.90) with the germline specific genes *glh-1* (Gruidl et al., 1996) and *mei-2* (Srayko et al., 2000) (Fig. 4C). This resulted in the identification of 4025 genes in hermaphrodites and 3051 genes in males (Table S3). As a validation of our approach, we found that most of these genes overlap with germline expressed genes that were identified by RNA sequencing of dissected gonads (94% for both the male and hermaphrodite germline genes) (Fig. 4D) (Ortiz et al., 2014). When the hermaphrodite and male data were combined, the total number of germline pattern genes was 4348. These comprise most germline specific genes that have been studied to date, including regulators of germ cell proliferation, meiosis and gamete production.

In addition to the 2728 germline pattern genes that were shared between the two sexes, there were 1297 genes that were only detected in hermaphrodites and 323 genes that were only detected in males (Fig. 4D). Examination of the expression patterns of these non-overlapping genes showed, however, that most are expressed in both the hermaphrodite and male germline regions (Fig. S4A), with differences in gonad structure precluding their detection in both sexes. These results indicate that the majority of the germline pattern genes are expressed in both the male and hermaphrodite germline. To further investigate if there are sex-specific differences in the expression of the germline pattern genes, we examined differential gene expression in the pooled male and hermaphrodite datasets using DESeq2 (Love et al., 2014). We found that 1274 genes were >2 fold enriched in hermaphrodites (adjusted p value < 0.1) and 3895 genes in males, which corresponds well with bulk RNA sequencing data (Kim et al., 2016). We found that 789 of the germline pattern genes were >2 fold enriched in hermaphrodites, 211 (27%) of which were only identified in the hermaphrodite expression maps (Fig. 4D, Table S3). Moreover, 330 of the germline pattern genes overlapped with a dataset of genes upregulated in feminized germlines Ortiz et al., 2014), consistent with a specific function of these genes in the oogenic germline. Finally, we found that 10 of the germline pattern genes were male enriched (>10 fold higher expression) (Table S3), and that these genes were only identified in the male expression maps. Among these was *fog-3*, which encodes a key regulator of spermatogenesis (Ellis and Kimble, 1995; Ortiz et al., 2014). We conclude that in addition to the majority of germline genes that are shared between males and hermaphrodites, there are also important differences in sex-specific expression. These observations are consistent with previous studies on dissected germlines (Ortiz et al., 2014), but our data and approach has the advantage that it can directly identify germline specific genes without the need to compare to somatic data.

We used a similar approach to examine sex-specific gene expression differences between male and hermaphrodite sperm. In hermaphrodites, sperm is produced for a brief period at the end of larval development and is stored in the spermatheca to take part in fertilization once the animal switches to oocyte production (Ellis and Stanfield, 2014; Nishimura and L’Hernault, 2010). Males produce sperm throughout adulthood and can mate with hermaphrodites to produce cross progeny. Interestingly, male sperm has an advantage over hermaphrodite sperm through a still poorly understood sperm competition mechanism (Ellis and Stanfield, 2014; Kulkarni et al., 2012; Singson et al., 1999). In males, initiation of spermatogenesis is marked by expression of *spe-44* (Kulkarni et al., 2012), while *spe-9* is expressed in developing spermatids (Putiri et al., 2004). We found that these genes showed a clear spatial separation in the male expression maps, corresponding to the linear assembly line like structure of the male gonad (Fig. 2F). To identify sperm specific genes in both hermaphrodites and males, we used the spermatid specific genes *spe-4* and *msp-3* as anchor genes (Arduengo et al., 1998; Kosinski et al., 2005). *spe-4* and *msp-3* were expressed in two sharp peaks in hermaphrodites, and in a single broad peak in males (Fig. 4E, S4A, B). We identified 1560 genes in males, and 1269 genes in hermaphrodites (1602 genes combined) that were co-expressed with either sperm marker (Fig. 4F, Table S3). As discussed above, these comprise the majority of the previously identified sperm specific genes and show 94% overlap with genes expressed in masculinized gonads (Ortiz et al., 2014). We found that the similarity to isolated mature male sperm (Ma et al., 2014) was lower (68%), which may reflect a difference in gene expression between the immature spermatids of young adults and the mature sperm of older adult males. Interestingly, while most (97%) of the sperm pattern genes identified in hermaphrodites overlapped with those in males, there were 333 genes that were only identified in males, 217 of which were >10 fold enriched in the male transcriptome (Fig. 4F, Table S3). We propose that these are male specific sperm genes that may contribute to the functional and morphological differences that enable male sperm cells to outcompete hermaphrodite sperm.

Taken together, these results show that using spatial transcriptomics, we can detect sex-specific differences in gene expression in sexually-dimorphic structures, such as the germline and sperm, and provide new insights into male and hermaphrodite specific aspects of reproductive biology.

### Spatial and sex-specific gene expression analysis identifies male specific genes in neurons and in the reproductive system

Genome-wide expression studies in *C. elegans* have mostly focused on the hermaphrodite, while the male transcriptome has remained relatively understudied (Kim et al., 2016). Motivated by the identification of male sperm genes, we therefore set out to characterize male specific genes that are expressed in cells required for mating and fertility. The successful mating of a male with a hermaphrodite depends on specific adaptations of the male nervous system and tail. Neurons such as the male specific CEM and MCM neurons in the head as well as mechanosensory and chemosensory neurons in the tail are required for detecting the hermaphrodite and locating the vulva (Sammut et al., 2015; Wang et al., 2014), while coupling to the vulva is mediated through the specialized structure of the male tail (Liu and Sternberg, 1995). Furthermore, sperm storage and ejaculation is dependent on parts of the male reproductive tract (including the seminal vesicle and vas deferens), which are also responsible for producing seminal fluid and the activation of spermatids prior to injection into the uterus of the hermaphrodite (Ellis and Stanfield, 2014; Kim et al., 2016; Palopoli et al., 2008; Smith and Stanfield, 2011).

The two pairs of CEM neurons (CEMV and CEMD) are located at a similar AP position in the head. The CEM neurons express genes related to the polycystic kidney disease *(pkd-2*) pathway, and share this profile with the HOB and RnB ray neurons in the tail (Barr and Sternberg, 1999; Wang et al., 2015; Wang et al., 2014). We used one of these genes, *cwp-1* (Fig. 1F), as an anchor gene to identify co-expressed genes in the head region (Fig. 5A, Table S3). Since the CEM neurons are only present in males, genes specific to the sections containing the CEM neurons are expected to be enriched in the male transcriptome. We found that 42 (24%) of the 173 genes co-expressed with *cwp-1* showed a >5 fold higher expression in males (Fig. 5B, Table S3). These included known CEM specific genes such as *cwp-2, pkd-2* and *trf-1*. Furthermore, 25 of the 42 male enriched genes were also found by bulk RNA sequencing of isolated extracellular vesicle releasing neurons (EVN), a subset of 27 different neurons that includes the CEM, HOB and RnB neurons (Wang et al., 2015). To further validate our results, we generated transcriptional reporters for *trf-1* and the uncharacterized gene F49C5.12. As expected, both genes were specifically expressed in the CEM neurons in the male head region (Fig. 5C, S5A). We conclude that combining pattern similarity with sex-specific gene expression is a powerful approach to identify gene expression in individual male neurons.

**Figure 5.**
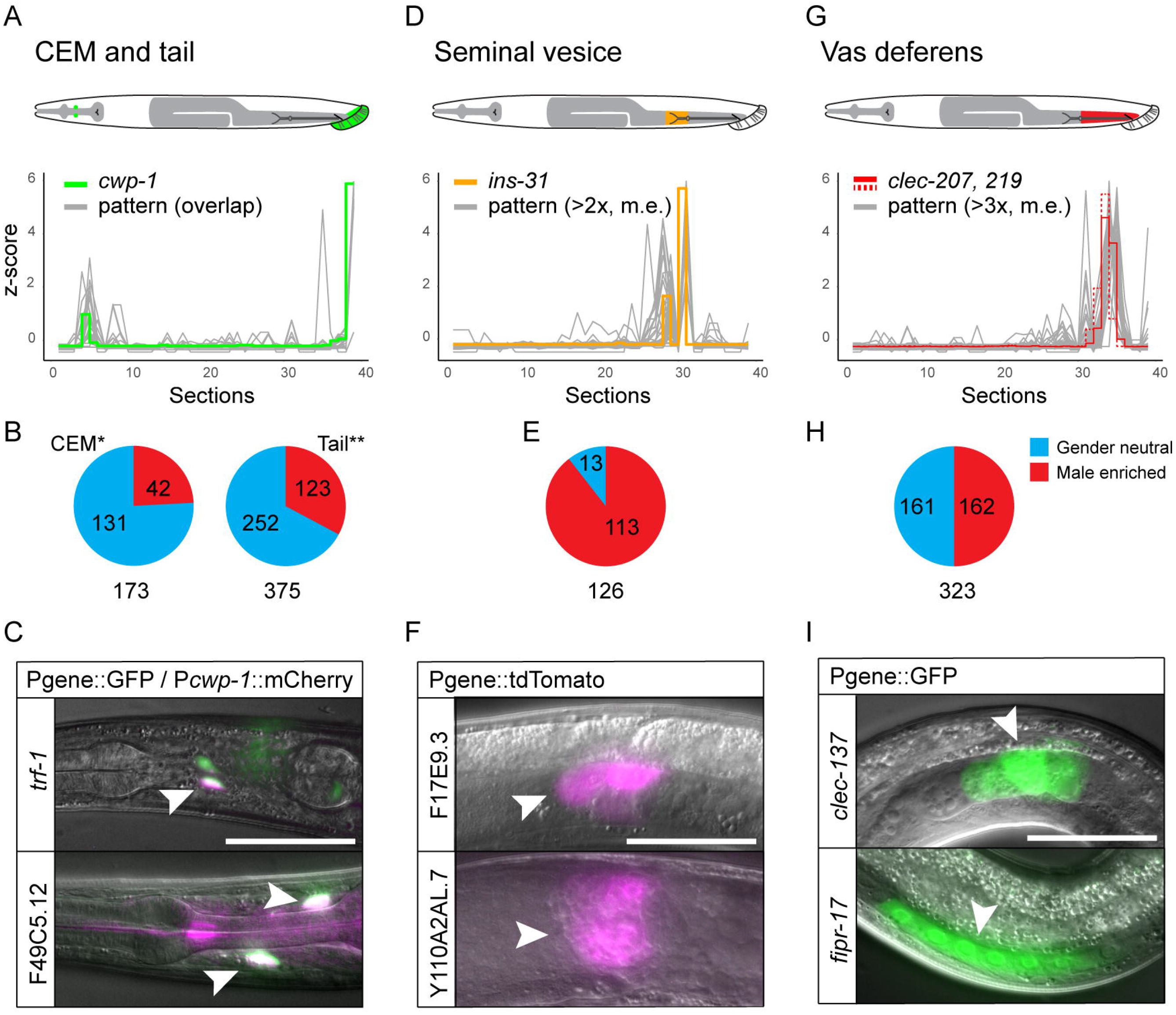
Identification of male specific genes expressed in male specific neurons and the reproductive tract. (**A**) Identification of genes expressed in the male CEM neurons and the HOB and RnB neurons in the tail. The anchor gene *cwp-1* is shown in green. Genes detected (r > 0.90) in both the CEM neurons and the tail (Table S3) are shown in grey. (**B**) Genes detected in the CEM neurons using *cwp-1* (*) and tail neurons using *cwp-1* and *flp-25* (**) as anchor points. The number of genes >5-fold enriched in males is shown in red. (**C**) Expression of transcriptional reporters of the CEM pattern genes *trf-1* and F49C5.12 in the CEM neurons (arrow heads). Scale bar is 25 μm. (**D**) Identification of genes expressed in the seminal vesicle. The anchor gene *ins-31* is shown in orange. Male enriched genes detected (r > 0.90) in at least 2 datasets are shown in grey. (**E**) 126 seminal vesicle pattern genes were identified, 113 of which are >5-fold enriched in the male transcriptome. (**F**) Validation of the expression of 2 seminal vesicle pattern genes (seminal vesicle indicated by arrow head). (**G**) Identification of genes expressed in the vas deferens. The anchor gene *clec-207* (solid line) and *clec-219* (dashed line) are shown in red. Male enriched genes detected (r > 0.90) in at least 3 datasets are shown in grey. (**H**) 323 vas deferens pattern genes were identified, 162 of which are >5-fold enriched in the male transcriptome. (**I**) Validation of the expression of 2 vas deferens pattern genes (arrow heads point to the vas deferens). For all reporters, at least 2 independent transgenic strains per gene were analysed.

Encouraged by these results, we set out to identify genes that are specifically expressed in the male tail, the seminal vesicle and the vas deferens. For the male tail, we selected *cwp-1* (for its expression in the HOB and RnB neurons) and *flp-25*, a neuropeptide gene that showed a specific peak of expression at the posterior end of our expression maps. We found 375 genes to be co-expressed with these marker genes, 123 of which were >5 fold enriched in males (Fig. 5B, Table S3). 26 of these genes overlapped with the CEM specific pattern genes, which is consistent with the shared EVN expression profile of these neurons (Wang et al., 2015).

To identify seminal vesicle specific genes, we used the previously identified marker gene *ins-31* as an anchor gene (Fig. 5D) (Kim et al., 2016). We found 126 genes with a similar peak pattern, the majority (90%) of which were male specific (>10 fold higher expression) (Fig. 5E, Table S3). Among these is *try-5*, a known seminal vesicle specific gene that is required for sperm activation (Smith and Stanfield, 2011). To further validate these results, we generated transcriptional reporters for two of the uncharacterized genes (F17E9.3 and Y110A2AL.7) and confirmed that both are specifically expressed in the seminal vesicle (Fig. 5F, S5B).

For the vas deferens, we used the markers *clec-209* and *clec-219* (Kim et al., 2016). We found 323 co-expressed genes, 162 of which were male specific (>10 fold higher expression) (Fig. 5G, H, Table S3). Again, we picked genes *(fipr-17, clec-137* and C09G12.5) for validation and found that all were specifically expressed in the vas deferens (Fig. 5I, S5C). Interestingly, a quarter of the vas deferens genes encode C-type lectins, a class of proteins that bind carbohydrates in a calcium dependent manner (Drickamer and Fadden, 2002). Together, these results provide unique insight into sex-specific gene expression in male neurons and the male reproductive system.

### Identification of novel male mating and fertility genes

The successful identification of genes specific to male neurons, the male tail, and the male reproductive tract prompted us to examine the function of the identified genes in mating and fertility. We measured mating efficiency as the ability of males to sire progeny. For this, we crossed RNAi treated males with *dpy-5* hermaphrodites. *dpy-5* mutants have an easily recognizable, recessive Dpy phenotype, enabling us to distinguish self-progeny (Dpy) from heterozygous cross-progeny (wild type). We depleted 30 male tail and CEM specific genes, 34 seminal vesicle genes and 57 vas deferens genes (Table S4) and found 10 genes that induced a significant decrease in cross-progeny (Table 1). Among these are 5 uncharacterized genes that encode small (62 - 133 amino acid) proteins, 2 genes that encode C-type lectins and 3 genes that have mammalian homologs. These include *clc-4*, which encodes a Claudin protein, *frm-8*, which encodes a FERM domain containing protein and *grd-4*, which encodes a Hedgehog-like protein. Interestingly, the uncharacterized proteins and the two C-type lectins are predicted to be secreted, indicating that they may represent novel components of the seminal fluid that are required for male fertility. Taken together, these results demonstrate that combining pattern-based gene identification with RNAi screens is a powerful new approach to identify novel tissue-specific gene functions.

**Table 1.**
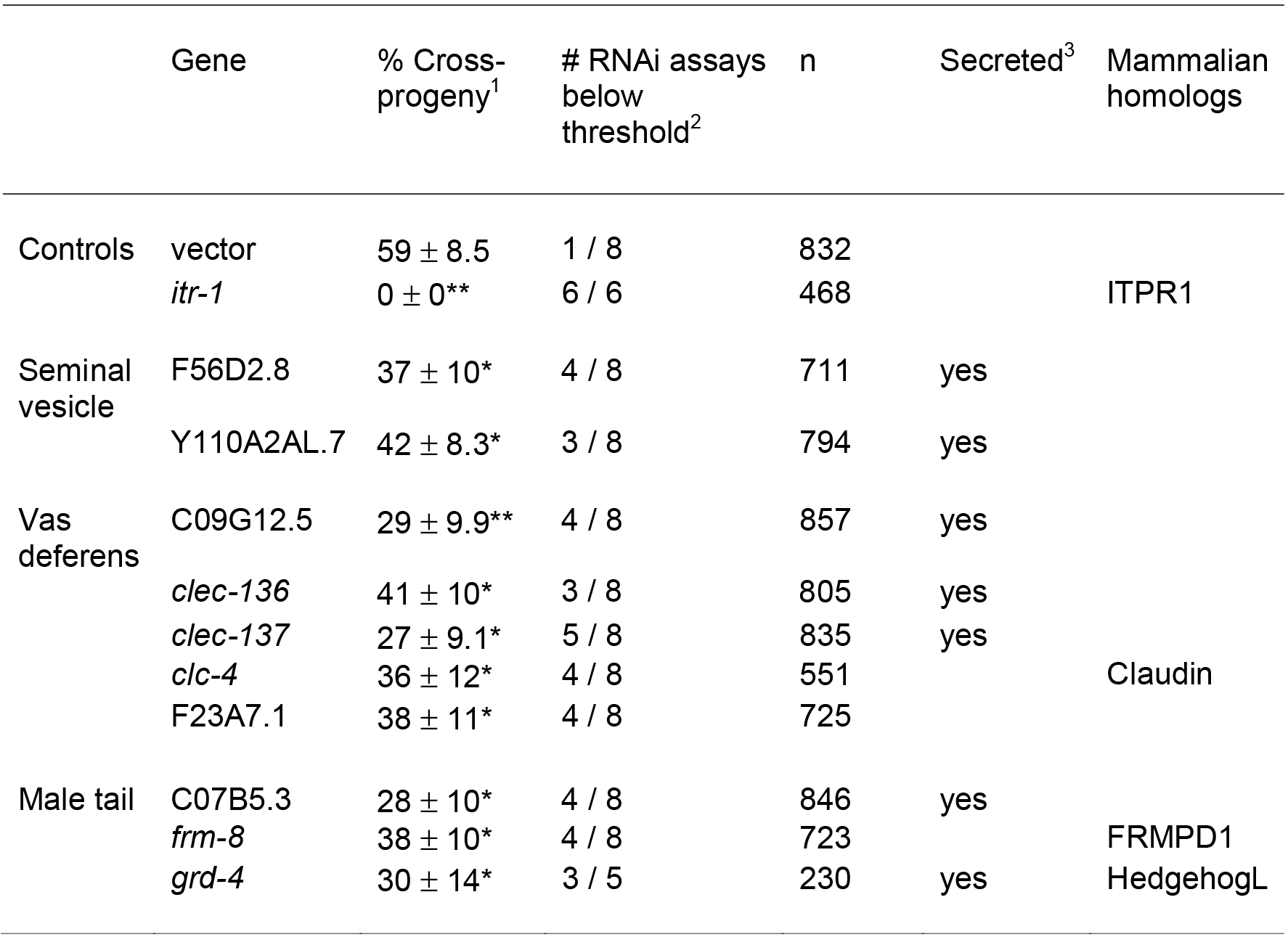
RNAi screen identifies genes required for male mating and fertility See supplementary methods for a description of the RNAi screening procedure. The inositol triphosphate receptor gene *itr-1* - which is required for turning behavior, spicule insertion and sperm transfer (Gower et al., 2005) - was used as a positive control. ^1^Percentage of crossprogeny (mean + SEM, *p < 0.05, **p < 0.01 compared to empty vector control, Student’s t-test). ^2^Number of RNAi assays showing a percentage of cross-progeny below a threshold of 35% (mean minus one standard deviation of the empty vector control). n, total number of animals scored. ^3^Genes that are predicted to encode secreted proteins (presence of signal sequence but no transmembrane domains, using SignalIP 4.1 server and TMHMM v2.0 server).

## Discussion

### Genome-wide expression maps of *C. elegans*: an expression pattern resource and a tool for tissue-specific gene identification

We used RNA tomography to generate the first genome-wide gene expression maps of *C. elegans* males and hermaphrodites. Despite the low amount of input material (with a sectioning width of 20 μm, sections outside of the germline contain only about 20-30 cells), this resulted in high complexity data, with the majority of young adult stage genes detected in our datasets. Examination of cell specific marker genes confirmed the high spatial resolution of the expression maps and demonstrated that genes expressed in small numbers of cells, such as specific neurons, can be readily detected. Since the anatomy of *C. elegans* is largely invariant (Sulston et al., 1980; Sulston and Horvitz, 1977), sensitivity could be further increased by aligning and pooling datasets. These pooled expression maps, which were generated by unbiased transcriptome-wide clustering as well as by manual alignment, are especially useful for analyzing lowly expressed genes.

In addition to examining expression patterns on a gene-by-gene basis, a powerful application of the expression maps is the identification of tissue-specific genes. We developed a computational approach that identifies co-expressed genes based on similarity in expression patterns and showed that this can be used in combination with marker genes to identify tissue-specific genes. An advantage of this approach over RNA sequencing of isolated cells or tissues is that it focuses on genes that are specific to the cell or tissue, while ubiquitously expressed genes are filtered out. We have identified genes that are specifically expressed in major organs and cell types of males and hermaphrodites, providing important new insight into tissue-specific gene expression profiles.

The expression maps and an interactive search tool for finding co-expressed genes are accessible on the project website. Both resources provide valuable information on the tissue-specificity and thus possible function of genes of interest, and are a starting point for further analysis using microscopy based techniques such as single molecule fluorescent *in situ* hybridization (smFISH) or fluorescently-tagged reporter transgenes.

### Gene expression maps as a tool to study sex-specific gene expression differences in male and hermaphrodite tissues

As an application of our approach to identify tissue-specific genes, we focused on sex-specific gene expression differences in the germline, sperm, and specific somatic cells of the male. Using meiotic markers in the germline as anchor genes, we identified over 4000 genes with a germline specific expression pattern. Among these are the majority of germline genes that have been described to date. We found that most germline pattern genes are shared between the two sexes, but that there are also clear sex-specific differences in expression levels that are consistent with the distinct spermatogenic and oogenic functions of the male and hermaphrodite germline. These results are in agreement with previous studies on dissected gonads (Ortiz et al., 2014), but are enriched for genes that are specific to the germline. Interestingly, we found that the difference between male and hermaphrodite sperm genes was more pronounced than what we observed for the germline. Thus, while most hermaphrodite sperm genes overlapped with male sperm genes, there were over 200 sperm pattern genes that were male specific. Male sperm has the ability to displace hermaphrodite sperm from the spermatheca to ensure the effective production of cross-progeny (Ellis and Stanfield, 2014; LaMunyon and Ward, 1998; Singson et al., 1999). This sperm competition mechanism is still poorly understood, but it is clear that the larger size and increased motility of male sperm plays a key role in this process. We suggest that the male specific sperm genes that we have identified may contribute to these morphological and functional differences.

We adapted this approach to identify genes expressed in specific somatic cells of the male. To extract the male specific genes from genes that are similarly expressed between the two sexes, we combined pattern similarity search with sex-specific differential gene expression analysis. This proved to be an effective way to find genes that are specifically expressed in the male CEM and tail neurons, and the seminal vesicle and vas deferens regions of the male reproductive tract, as shown by the detection of previously identified marker genes for these tissues and validation with transgenic reporters. RNAi mediated depletion of these male specific genes identified 10 genes that are essential for male mating or fertility. Interestingly, we found that 5 of the 7 seminal vesicle and vas deferens genes with a male fertility phenotype encode small, secreted proteins. One of the functions of the seminal vesicle and vas deferens is the production of seminal fluid. The seminal fluid is required for the ejaculation of sperm cells during copulation, but also plays an important role in sperm activation and potentially in sperm competition (Ellis and Stanfield, 2014; Kim et al., 2016; Palopoli et al., 2008; Smith and Stanfield, 2011). We propose that the 5 secreted proteins that we have discovered are novel components of the seminal fluid that play an essential role in male fertility. These results illustrate the power of combining pattern similarity analysis with RNAi screens to identify novel tissue-specific gene functions.

### Applications of spatially resolved transcriptomics in *C. elegans* and other nematodes

The expression maps that we have generated give a detailed overview of gene expression patterns in young adult males and hermaphrodites. A valuable addition will be the generation of expression maps of the different larval stages, which will provide insight into gene expression pattern changes during post-embryonic development. Importantly, since our work shows that expression maps from different animals can be aligned, RNA tomography can also be used as a powerful tool for comparative studies. For example, gene expression patterns can be compared between wild type and mutant animals, or between the laboratory strain Bristol N2 and strains isolated from the wild. Since nematodes share a common body plan, such comparisons can be extended to other nematode species to analyze spatially resolved gene expression differences in an evolutionary context. Such studies will be especially useful for the functional annotation of the many uncharacterized genes in nematode genomes.

## Author contributions

A.E., J.P.J, A.O. and H.C.K. conceived the study. A.E. and M.B. performed the cryo-section and mRNA sequencing. A.E. and M.B. generated the reporter lines and performed the RNAi screen. A.V., B.S., A.E. and E.B. performed the data analysis. E.B. made the website. A.E., A.V. and H.C.K. wrote the manuscript.

## Acknowledgements

We thank the Utrecht Sequencing Facility (USF) and Kay Wiebrands for technical help. This work is part of the research program (14NOISE01) of the Foundation for Fundamental Research on Matter (FOM), which is financially supported by the Netherlands Organization for Scientific Research (NWO). Some strains were provided by the CGC, which is funded by the NIH Office of Research Infrastructure Programs (P40 OD010440). We are grateful to Steve l’Hernault and members of the Korswagen and van Oudenaarden groups for helpful discussions, and to Maya Sen for critically reading the manuscript.

## Supplementary Figure legends

**Supplementary Figure 1 (related to Figure 1).**
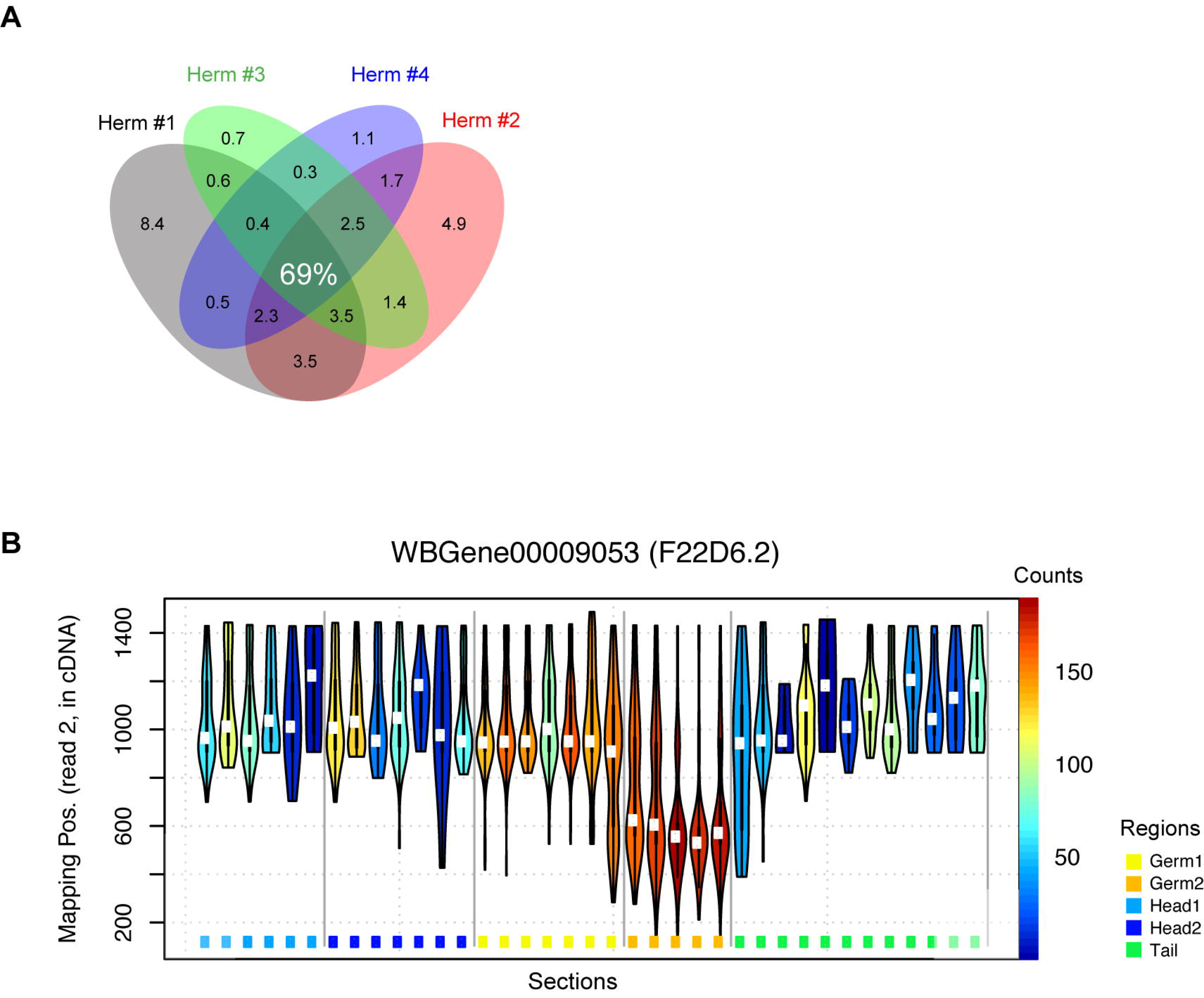
(**A**) Venn diagram of genes with >10 transcripts in at least 1 section in the 4 hermaphrodite expression maps. 69% of genes are detected in each of the 4 datasets. (**B**) Example of spatial differences in 3’ isoform usage of the gene F22D6.2 in Male #3. Violin plots show the distribution of mapping positions per section along the animal. The colors of the violins show the expression level (UMI count) per section (color scale on the right hand side). Colorful dots at the bottom and the grey vertical lines indicate the anatomical regions, and their boundaries as determined in Fig 3.

**Supplementary Figure 2 (related to Figure 2).**
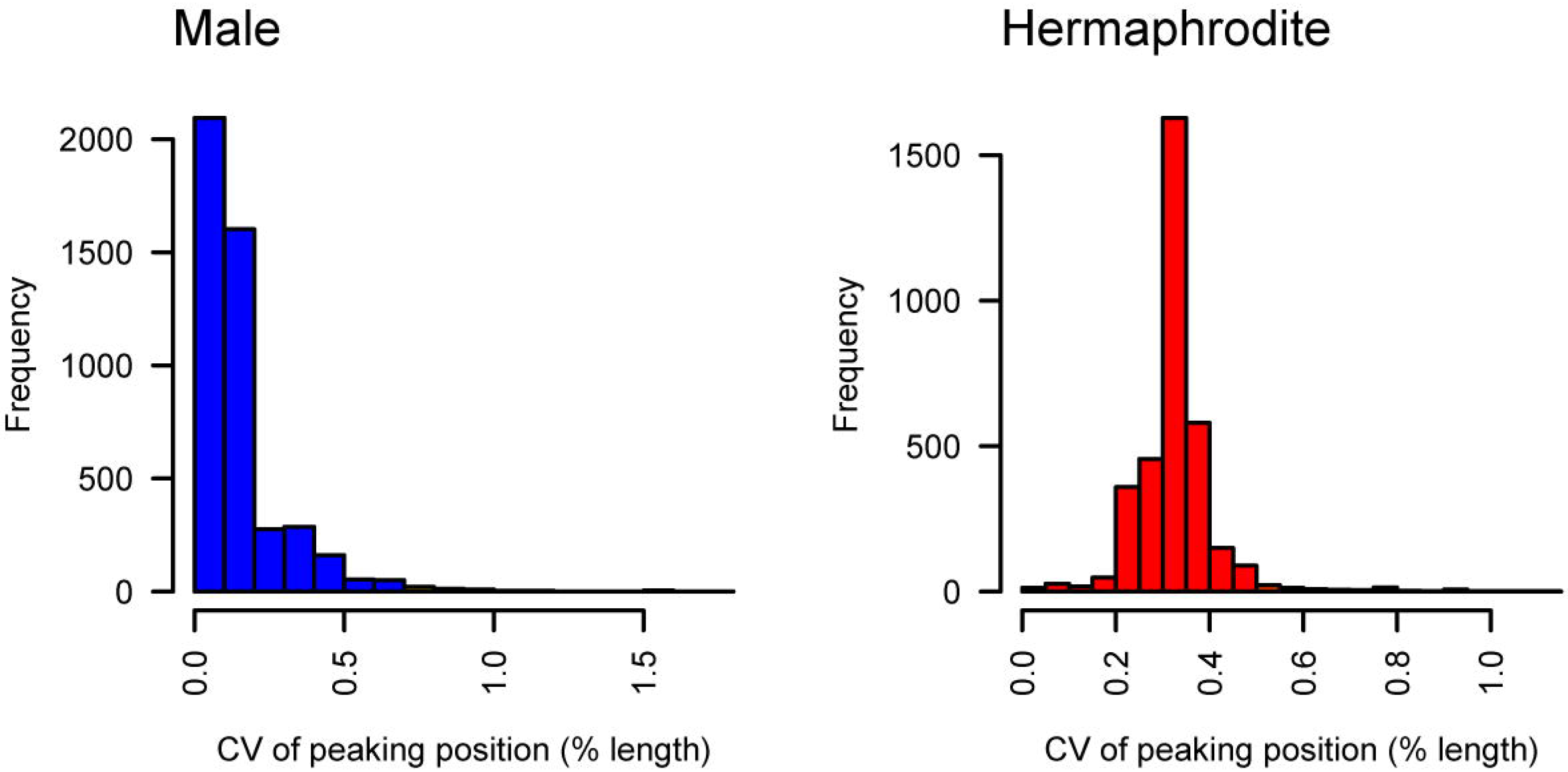
Coefficient of variation of relative peak position of genes displayed in Figure 2. Maximum expression position is calculated for each gene, and is expressed as percentage distance from the head in each animal. Coefficient of variation (CV) of relative position was calculated over all four animals of the same sex.

**Supplementary Figure 3 (related to Figure 3).**
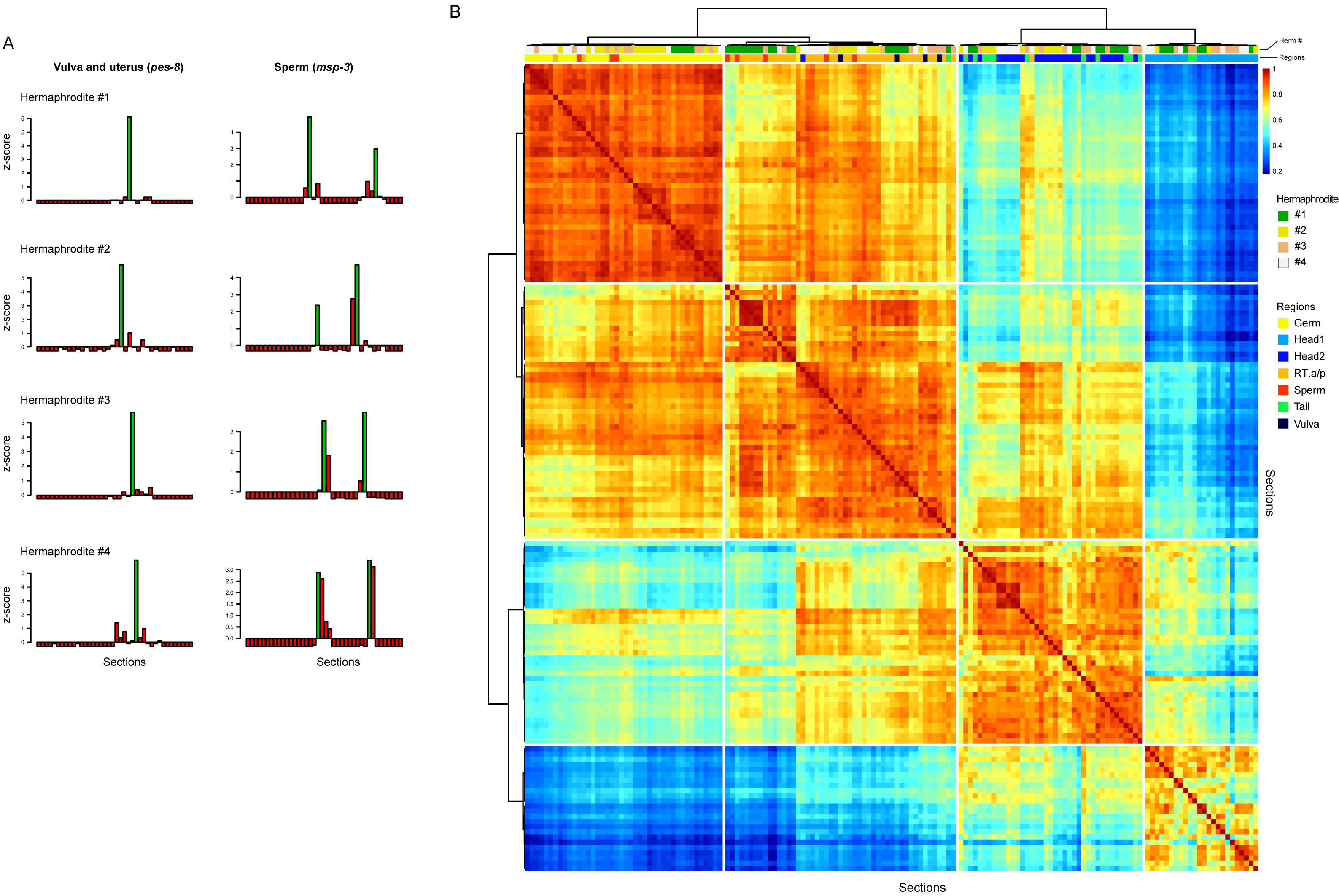
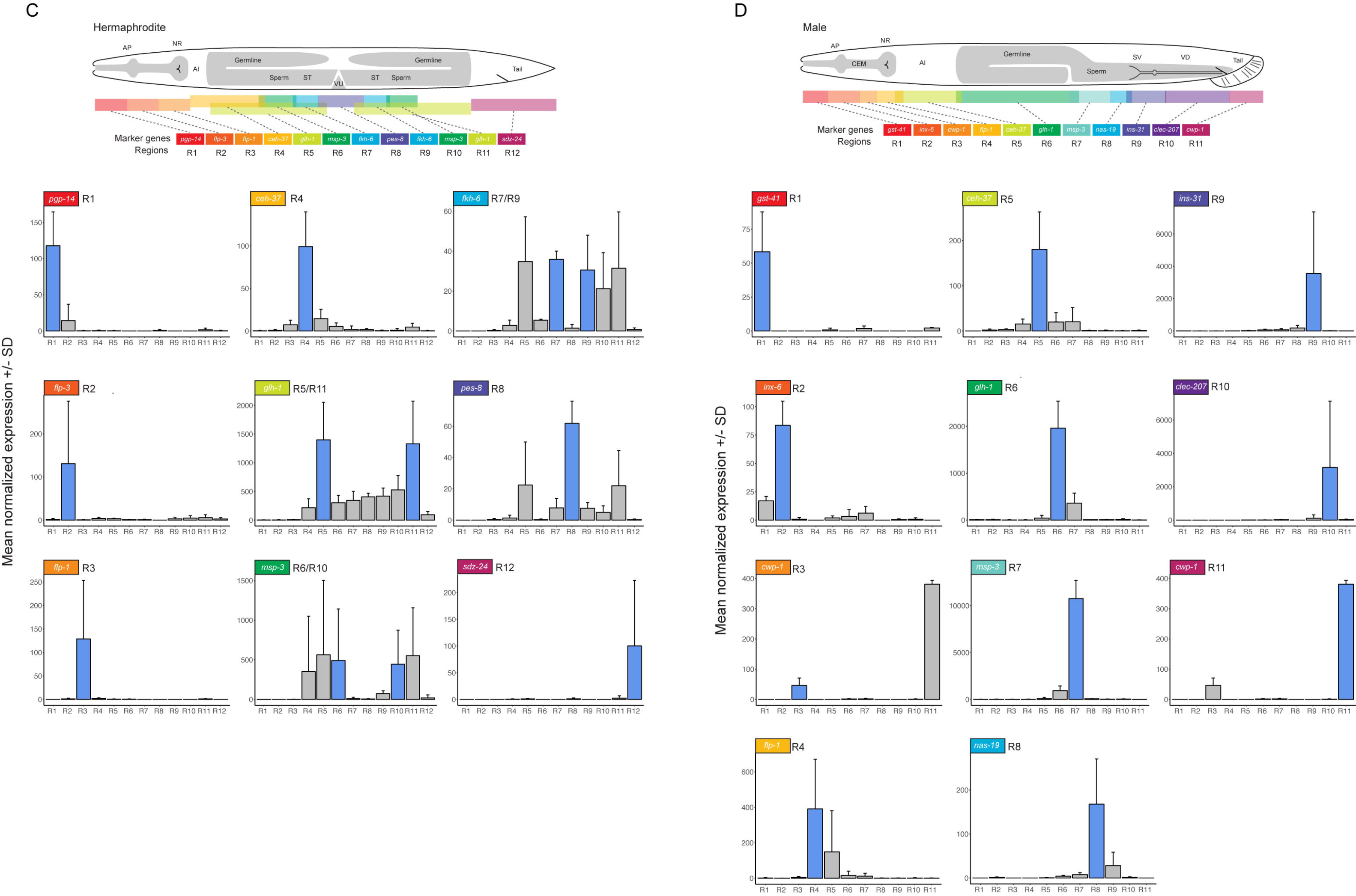
(**A**) Position of vulva/uterus *(pes-8)* and sperm *(msp-3)* expression peaks in the 4 hermaphrodite expression maps. The expression maxima were used to define reference points in the symmetrical hermaphrodite gonads to separate anterior and posterior parts of regions determined by hierarchical clustering. (**B**) Multi animal correlation for hermaphrodites, counterpart of Figure 3C. As in males, major regions cluster by anatomical identity rather than by sample of origin when all sections from all four hermaphrodites are clustered together. (**C**, **D**) Expression of the marker genes used for manual alignment in the merged datasets. Expression is mean normalized transcript count ± standard deviation. AP anterior pharynx region, NR nerve ring, AI, anterior intestine region, ST spermatheca, VU vulva and uterus region, SV seminal vesicle, VD vas deferens.

**Supplementary Figure 4 (related to Figure 4).**
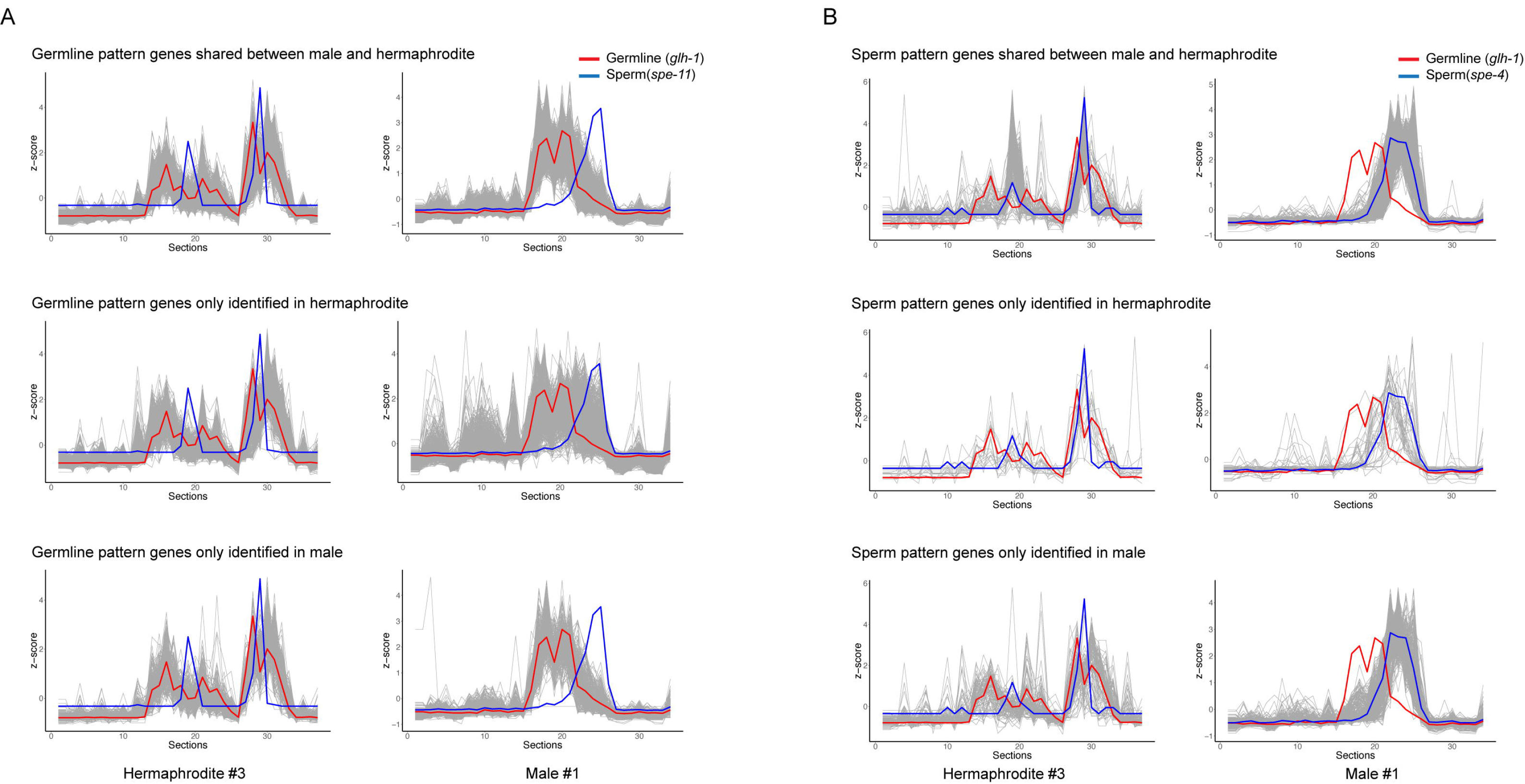
(**A**) Expression patterns of germline pattern genes (identified at r > 0.90) in males and hermaphrodites. Most germline genes only identified in hermaphrodites also show expression peaks in the male gonad. (**B**) Expression patterns of sperm pattern genes (identified at r > 0.90) in males and hermaphrodites.

**Supplementary Figure 5 (related to Figure 5).**
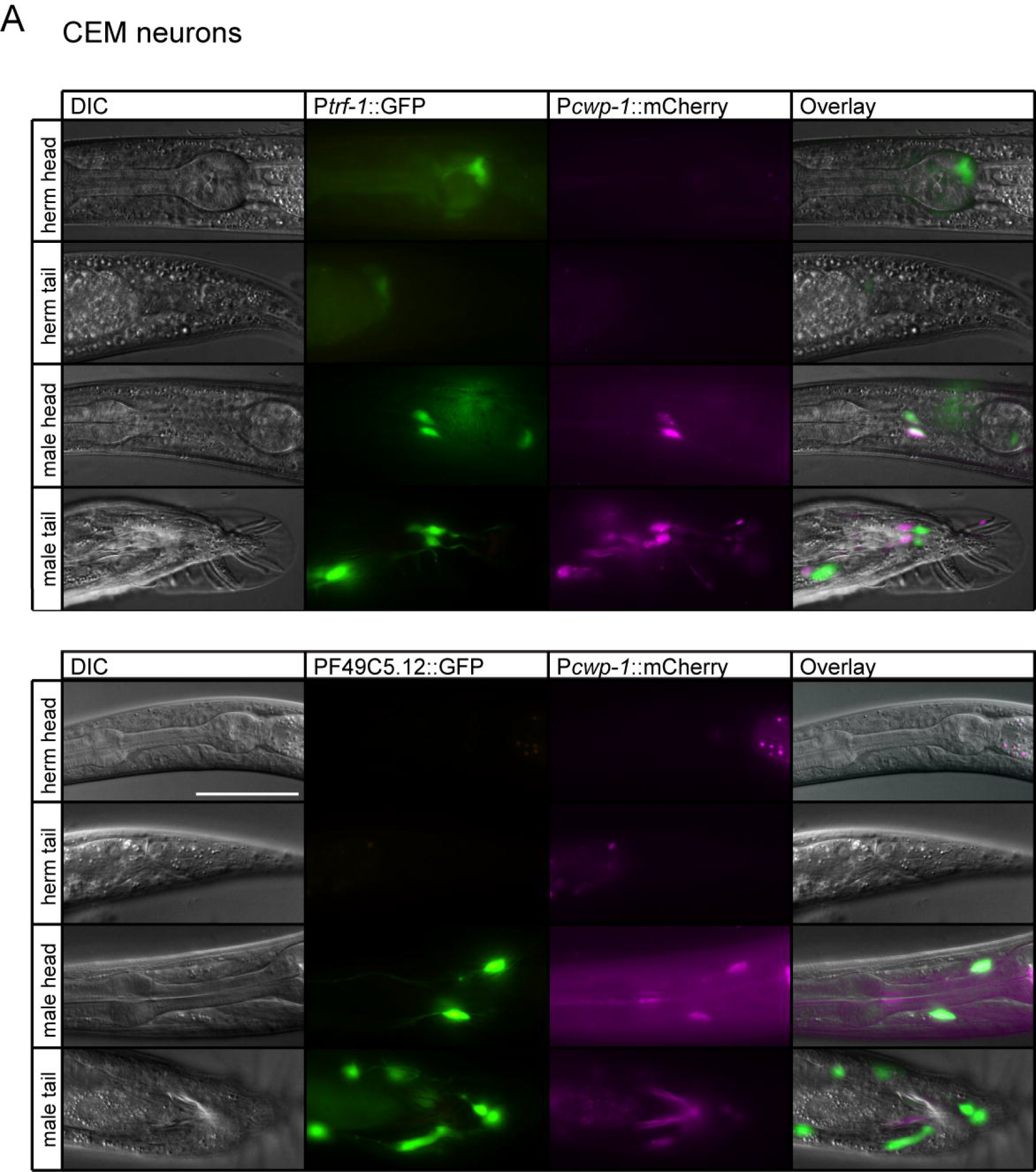
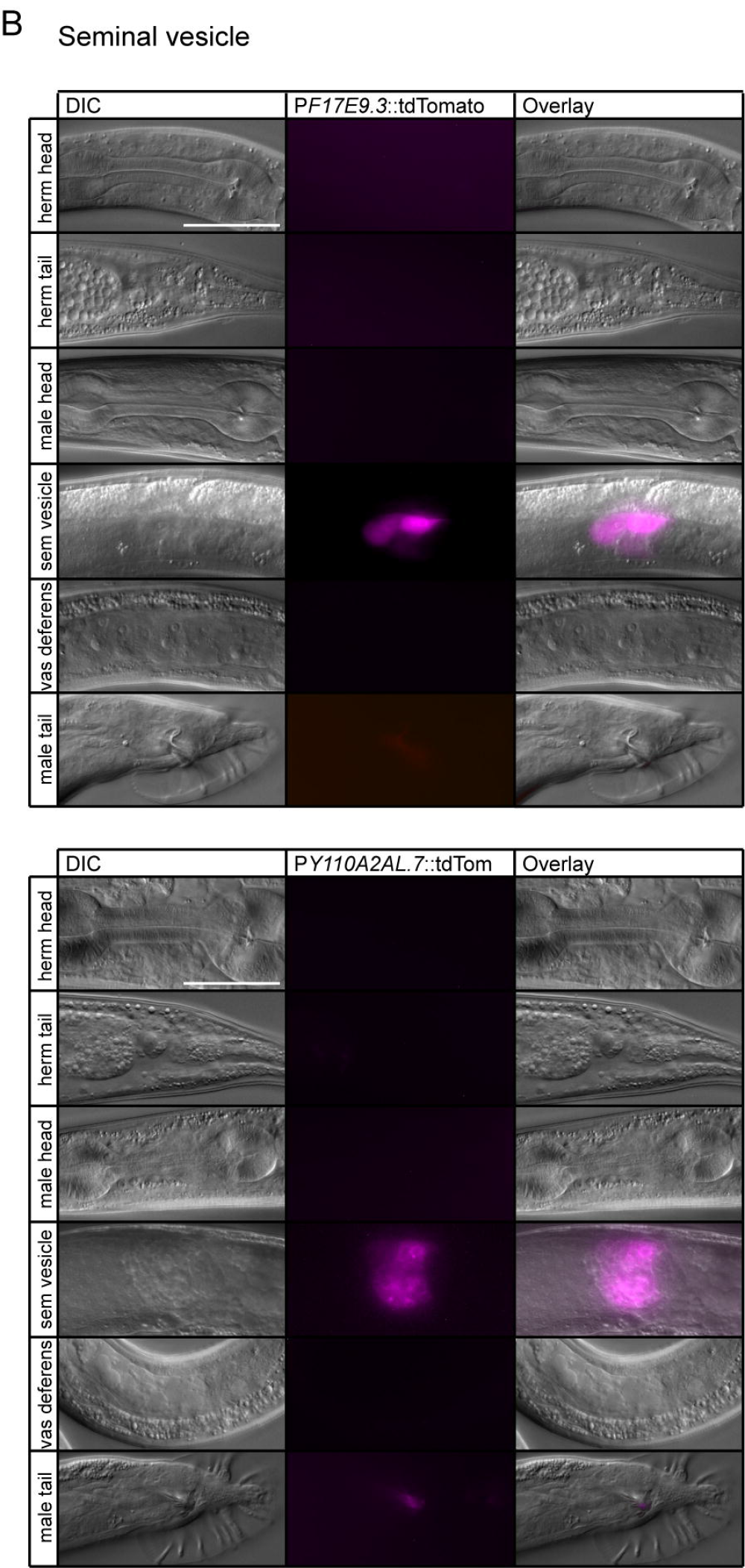
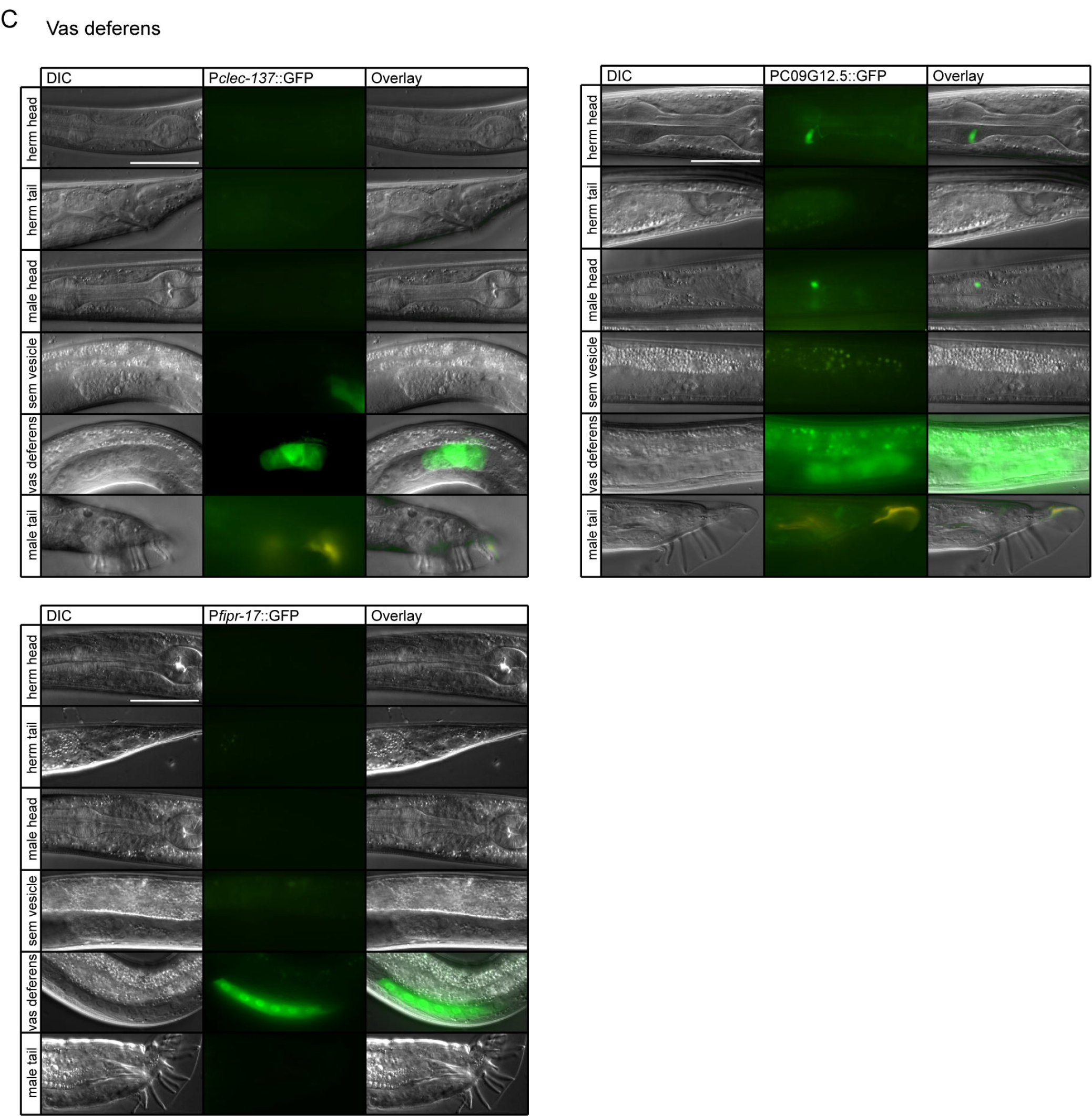
Validation of male specific pattern genes expressed in male specific neurons and the reproductive system. (**A**) Expression of *Ptrf-* 1::GFP and PF49C5.12::GFP reporter strains in the male head and tail region. In the male head the expression pattern overlaps that of *Pcwp-1* ::mCherry, confirming expression in the CEM neurons. In the tail the expression pattern only partially overlaps with *Pcwp-* 1::mCherry. No expression of F49C5.12 was observed in hermaphrodites. Scale bar is 25 μm. (**B**) Expression of the seminal vesicle pattern gene reporters PF17E9.3::tdTomato and PY110A2AL.7::tdTomato in the mid-tail region, which was confirmed to be the seminal vesicle by DIC imaging. No expression was observed in hermaphrodites. (**C**) Expression of the vas deferens pattern gene reporters P*clec-137*::GFP, P*fipr-17*::GFP and PC09G12.5::GFP in the posterior part of the male tail, which was confirmed to be the vas deferens by DIC imaging. No expression was observed in hermaphrodites, except for an unidentified cell in the head region for C09G12.5. For all reporters, at least 2 independent transgenic strains per gene were analyzed.

## Materials and Methods

### *C. elegans* strains and culture

Unless noted otherwise, *C. elegans* strains were cultured at 20°C using standard conditions (Lewis and Fleming, 1995). For RNA tomography, young adult Bristol N2 males and hermaphrodites, or young adult *glp-1(q231)* mutants (grown at the restrictive temperature of 25°C) were used. Other mutant alleles were *dpy-5(e61)* and *him-5(e1490)*.

### *C. elegans* reporter constructs and transgenesis

Transcriptional reporters were generated by combining approximately 1500 bp of upstream promoter sequence with fluorescent reporter genes. For *cwp-1*, F17E9.3 and Y110A2AL.7, PCR amplified promoter fragments were combined with mCherry or tdTomato and the *unc-54* 3’ UTR through Gateway cloning. For all other genes, the promoter fragment was fused to GFP and the *unc-54* 3’ UTR through fusion PCR (Hobert, 2002). All reporter constructs and PCR products were injected into *him-5(e1490)* at 30-50 ng/μl with the co-injection marker *rol-6(su1006)* (pRF4) at 30 ng/μl and pBluescriptII to a total concentration of 150 ng/μl. Table S5 lists the transgenic reporter strains and DNA oligonucleotides used in this study. For epifluorescence and DIC imaging animals were mounted on 2% agarose pads containing 10 mM sodium azide. Images were acquired using a Zeiss Axioscope microscope equipped with a Zeiss Axiocam digital camera. Figures were prepared using ImageJ software.

### RNAi screen

Bacterial clones for feeding RNAi of 121 candidate genes and the positive control *itr-1* (Gower et al., 2005) were selected from the Vidal and Ahringer libraries (Kamath et al., 2003; Rual et al., 2004) (Table S4). To assay male fertility, four *him-5(e1490)* hermaphrodites were grown on RNAi bacteria at 15°C for 5 days. For the initial screen, performed in duplicate, 6 adult male offspring were crossed with 3 L4 *dpy-5(e61)* mutants and kept at 20°C. Adult males were removed after 1 day, and after 3 days the number of self-progeny (Dpy) and cross-progeny (WT) was counted. Clones that scored positive were further tested 5-8 times with male/hermaphrodite ratios of 5:3, 2:2, and 1:1.

### Preparation of sequencing libraries

Live young adult males or hermaphrodites were oriented perpendicular to the sectioning axis, frozen and cryo-sectioned as described (Junker et al., 2014) with a sectioning width of 20 μm. mRNA extraction, barcoding, reverse transcription and in vitro transcription were performed as described (Junker et al., 2014) according to the CEL-seq protocol (Hashimshony et al., 2012) using the Message Amp II kit (Ambion). Illumina sequencing libraries were subsequently prepared according to the CEL-seq2 protocol (Hashimshony et al., 2016) using the SuperScript^®^ II Double-Stranded cDNA Synthesis Kit (Thermofisher), Agencourt AMPure XP beads (Beckman Coulter), and randomhexRT for converting aRNA to cDNA using random priming. The libraries were sequenced paired-end at 50 bp read length on an Illumina HiSeq 2500.

## Data analysis

### Read alignment (mapping)

We aligned the 50 base pair paired-end reads to the *C. elegans* references transcriptome, which was compiled from the *C. elegans* reference genome WS249. The transcriptome reference is available in fasta format on GitHub, under: http://bit.ly/Reference_C_elegans. For alignment, we used our custom wrapper MapAndGo2 (available on GitHub, https://bit.ly/MapAndGo_v2) around BWA MEM (Li and Durbin, 2010).

### Transcript counting

MapAndGo2 aligns read 2 against the forward strand of the reference transcriptome, and selects uniquely mapped reads (mapq ≥20 and no ‘XA’ or ‘SA’ BWA tags). Read 1 contains the sample-barcode, and the unique molecular identifier (UMIs) and it is used for transcript counting. First, uniquely mapped reads are assigned to samples using the sample barcodes. In the next step, duplicate reads are removed using UMIs. These reads originate from the same original mRNA molecule and are amplification duplicates. Such reads are defined by mapping to the same gene in the same cell and having the same UMI. Finally, we converted the number of observed UMIs into transcript counts based on random sampling with replacement (Grun et al., 2014).

### Analysis of count data

We analysed count data in R (v 3.3.2). Custom analysis scripts will be accessible upon acceptance at https://github.com/vertesy/C.elegans.Spatial.Transcriptomics. We used the R-packages: pheatmap, MarkdownReports, ggplot2 and corrr for visualization, DESeq2 for differential gene expression analysis, and a set of other custom scripts, which are available under https://github.com/vertesy. The mapping reference contained cosmid ID-s, which were converted first to WB-gene identifiers, then to gene names. Cosmid or Wormbase gene IDs were retained for genes where no matching Wormbase gene ID or gene name was found.

### Filtering genes and sections

To select robustly detected/quantified genes, we kept all genes expressed above 10 unique transcripts in at least one section for further analysis. This means that genes are selected on sufficient evidence (enough reads to robustly conclude), and not on “high expression” as measured by relative/normalized read counts. This is important, because many analyses are based on z-score transformed values, and sporadic gene expression (or detection) can yield high z-scores (~relative expression peaks). Sequencing libraries were then normalized to 10 million total transcripts per animal, to account for differences in sequencing depth. Next, the anterior and posterior ends of the expression maps were defined by the first or last two consecutive sections with ≥50 genes (with ≥20 normalized transcripts). Internal sections with <50 genes were removed (an average of 1.1 sections per animal).

### Alternative 3’ isoform mapping

CEL-seq is a 3’ sequencing protocol where transcripts are reverse transcribed from the 3’ end of the polyadenylated mRNA, using a poly-T primer. Read 1 is located directly at the beginning of the poly-A tail (3’ end of UTR), while the mapped read 2 lies ~400 bases upstream. This distance is determined by the fragmentation of the amplified RNA, after which the Illumina primers are ligated to the 5’ end of the fragmented transcripts, determining the start position of read 2. Although fragmentation is not deterministic, we reproducibly generated 300-600 bases long cDNA Illumina sequencing libraries in all 8 animals. We detected robust differences of mapping position of multiple genes along the A-P axis. These differences in many cases exceeded a kilo base, and changes corresponded to borders of the independently determined anatomic regions. The most notable differences in 3’ usage were detected between the soma/germline boundary and within different regions of the germline. To systematically identify these genes, we first extracted the mapping positions of uniquely mapped, UMI corrected reads from the sam files. We only considered genes identified as highly expressed in the rest of the analysis. We then calculated the median mapping position per gene, per section. To identify robust expression patterns, we selected genes with at least 10 UMI’s in 2/3 of the sections. Next, we identified genes with high positional variation, by selecting genes with interquartile range of median mapping positions above 100. To filter out genes with inconsistent, section-to-section variation in mapping positions, we calculated the autocorrelation of median mapping positions along the animal (lag of 1), and selected genes with a positive autocorrelation value. While higher autocorrelation values reproducibly identified region specific 3’ usage across the four males, it led to many false negatives in hermaphrodites because of their more complex anatomy. Therefore, we translated the question into a segmentation problem, which is a concept from copy number calling in DNA-sequencing. Segmentation algorithms identify spatially consistent signal differences. We provided the Bioconductor package DNAcopy with the median mapping positions and selected genes that contained at least 2 different segments at least 200 bases away, with the smaller segment of at least 3 consecutive sections. The resulting list was manually corrected for cases were differences were due to inconsistencies in gene models (Table S2). Genes with the most interesting patterns are presented in Supplementary dataset 1.

### Normalization and transformation

Datasets were normalized to 10 million reads per animal for most analyses. Sections within the boundaries of the animal were z-score transformed prior to normalization to 10 million transcripts. We provide the normalized and z-score transformed expression of all genes in supplementary dataset 1. For specific analyses (Heatmaps in Fig. 2A and B), transcript counts were converted to transcript per million (TPM) values per section, and then z-score transformed. We normalized each filtered, high quality dataset so that each section has 1 million transcripts, so that we can quantify and compare relative expression as transcript per million (TPM) values. Note that in CEL-seq only one 3’ fragment is sequenced, hence there is no need to normalize for mRNA length: it is only necessary for full length mRNA protocols, such as SMART-seq. z-score transformation after TPM normalization reduces technical slice-to-slice variation, but erases the ~5 fold germline-to-non-germline difference in RNA-content. Consequently it gives a more homogenous pattern within somatic and germline regions, but they only represent relative expression within each slice.

### Comparison to dataset Kim et al., 2016

A recent publication quantified sex-specific gene expression in *C. elegans*, sequencing mRNA in up to 600 animals per sample (Kim et al., 2016). We downloaded the normalized count data (Table S1) and we counted all detected genes in the M5 (young male) and in the H5 (young hermaphrodite) samples. Next, we similarly counted genes in our datasets (4 males and 4 hermaphrodites) and found 93% and 95.3% overlap between the hermaphrodite and male datasets, respectively.

### Gene Clustering

To visualize global gene expression patterns along the AP axis we clustered and displayed the transcript-per-million normalized, z-score transformed gene expression in all sections by hierarchical clustering using the pheatmap package with default parameters. (Fig 2A, B).

### Identification of anatomic regions

We calculated the pairwise Pearson correlation coefficients for all sections using all genes. Next, we sequentially broke down the animals into smaller and smaller regions using hierarchical clustering. Males are first separated into 3 clusters representing 2 germline-associated and a somatic section. We found that in male #1 we needed to cluster all germline sections together into 2 clusters to properly separate germ1 and germ2. In case of the hermaphrodites we made use of 2 marker genes. We used the single maximal expression of *pes-8* to define the vulva, which is the symmetry axis of the germline structure. We also identified the symmetric, sperm containing sections by the expression maxima of *msp-3* anterior and posterior of the vulva (Fig. S3A).

### Co-clustering of regions across animals

We show that the anatomic regions identified in individual animals are reproducible across different animals, and that they are not affected by batch or individual-to-individual variation. Therefore, we combined all datasets from the same sex and clustered all sections using Pearson correlation, as before. The upper bars in Fig. 3C and Fig. S3B indicate that the sections cluster by the regional identity determined in individual animals and do not cluster by sample of origin.

### Manually curated alignment of animals

Marker genes at specific locations along the AP axis were selected from genes displaying specific peak expression patterns. Normalized datasets were aligned and pooled based on z-scores of marker gene expression. In case a section showed overlapping marker gene expression (z-score > 1), the overlapping section was added to both regions. Sections with no marker gene expression (z-score < 1) were added to the closest matching region.

### Alignment and pooling of regions

After the identification and validation of anatomical regions, we added up transcript counts for all section either per clustering based regions or per manually curated regions. The manually curated datasets can be browsed at the project website (http://celegans.tomoseq.genomes. nl).

### Identification of spatially co-expressed genes

We implemented a similarity search algorithm which identifies all genes similar to a chosen gene of interest that is accessible on our website (http://celegans.tomoseq.genomes.nl). Similarity across sections is calculated using either Pearson or Spearman correlation or by Euclidean or Manhattan distance of gene expression. Either TPM-normalized or z-score normalized data is suggested to use, but any genes by sections (rows, columns) formatted expression data matrix can be fed to the function.

## Quantification and statistical analysis

For the RNAi screen, the number of animals scored (n), the mean and SEM and the statistical significance are reported in the Table legend. Data were regarded statistically significant when p < 0.05 by two-tailed Student’s t-test.

## Data resources

The RNA-seq data have been deposited at the Gene Expression Omnibus (GSE114723). The analyzed data set is available on the project web page (http://celegans.tomoseq.genomes.nl), including display of (1) z-score normalized gene expression, (2) merged, region specific gene expression and (3) spatially co-expressed genes.

## Supplementary Table legends

**Supplementary Table 1.** Overview of RNA sequencing data. Reads, total number of mapped sequencing reads. Transcripts, total number of transcripts based on UMI correction. Genes, total number of genes defined by ≥1 or ≥10 transcripts.

**Supplementary Table 2.** Alternative 3’ isoform mapping. The table summarizes how many times each of the 39 genes has been detected, in all, in hermaphrodite, or in male animals.

**Supplementary Table 3.** Tissue-specific pattern genes. Marker genes (reference genes) and datasets are indicated for each tissue. The threshold for similarity in expression pattern is a Pearson correlation value (r) of >0.90. For each gene, the cumulative r values for the male datasets (Sum R male), the hermaphrodite datasets (Sum R herm) and the total sum of r values over all datasets (Sum R total) are indicated. At a cumulative r value of >1.8, a gene is observed in at least two datasets. In case of the germline, sperm and the male reproductive tract, additional information is provided on male or hermaphrodite enriched genes. Known tissue-specific genes are indicated in green.

**Supplementary Table 4.** Male neuron and reproductive tract pattern genes tested in RNAi screen for male fertility defects.

**Supplementary Table 5.** Transgenic lines and DNA oligonucleotides that were generated and used in this study.

## Supplementary Datasets

**Supplementary Dataset 1** (Related to Figure S1B). Spatial differences in 3’ isoform usage. Violin plots show the distribution of mapping positions per gene, per section along the animal. The colors of the violins show the expression level (UMI count) per section (color scale on the right hand side). Colorful dots at the bottom and the grey vertical lines indicate the anatomical regions, and their boundaries as determined in Fig.3. Bottom right legend shows 3 basic statistics: The 33^rd^ quintiles of median UMI count (q33) is indicative of expression throughout the animal; Interquartile range of median mapping positions (IQR) indicates the variability in 3’ mapping, while the lag=1 autocorrelation (ACF) indicates spatial consistency of the mapping positions.

**Supplementary Dataset 2** (Related to Figure 3). Pairwise Pearson correlation heatmaps of the different male and hermaphrodite datasets.

